# Retinoschisin deficiency induces persistent aberrant waves of activity affecting neuroglial signaling in the retina

**DOI:** 10.1101/2021.08.26.457777

**Authors:** Cyril G. Eleftheriou, Carlo Corona, Shireen Khattak, Elena Ivanova, Paola Bianchimano, Yang Liu, Duo Sun, Rupesh Singh, Julia C. Batoki, Jason J. McAnany, Neal S. Peachey, Carmelo Romano, Botir T. Sagdullaev

**Affiliations:** Burke Neurological Institute at Weill Cornell Medicine, White Plains, NY 10605; Regeneron Pharmaceuticals, Inc. Tarrytown NY 10591; Cole Eye Institute, Cleveland Clinic, Cleveland OH 44195; Department of Ophthalmology, University of Illinois at Chicago, Chicago Illinois; Louis Stokes Cleveland VA Medical Center, Cleveland OH 44106; Department of Ophthalmology, Cleveland Clinic Lerner College of Medicine of Case Western Reserve University, Cleveland OH 44195

**Keywords:** XLRS, retinoschisis, retinal development, optophysiology, two-photon microscopy

## Abstract

Genetic disorders which present during development make treatment strategies particularly challenging because there is a need to disentangle primary pathophysiology from downstream dysfunction caused at key developmental stages. To provide a deeper insight into this question, we studied a mouse model of X-linked juvenile retinoschisis (XLRS), an early onset inherited condition caused by mutations in the *RS1* gene encoding retinoschisin (RS1) and characterized by cystic retinal lesions and early visual deficits. Using an unbiased approach in expressing the fast intracellular calcium indicator GCaMP6f in neuronal, glial, and vascular cells of the retina of mice lacking RS1, we found that initial cyst formation is paralleled by the appearance of aberrant spontaneous neuro-glial signals as early as postnatal day 13. These presented as glutamate-driven wavelets of neuronal activity and sporadic radial bursts of activity by Müller glia, spanning all retinal layers and disrupting light-induced signaling. This study highlights a critical role for RS1 in early retinal development with a potential to disrupt circuit formation to central targets. Additionally, it confers a functional role to RS1 beyond the scope of an adhesion molecule and identifies an early onset for dysfunction, a potential temporal target for therapeutic intervention and diagnosis.

**Significance Statement/ Highlights:**

- Photoreceptor inner segments express *Rs1* at P5, after which RS1 protein is detected in the inner segments by P9 and throughout the retina at later ages, with structural abnormalities observed by optical coherence tomography at P13 in *Rs1* mutant mouse models.
- Aberrant glutamate-driven wavelets identified by GCaMP6f-based analyses are a novel pathophysiological feature of RS1 deficient mice that emerge after maximal RS1 expression.
- Müller glia display abnormal radial glutamate-driven coordinated and sporadic bursts of activity in RS1-deficient mice.
- These data identify a novel pathophysiological feature of RS1-deficient mice and define a window where treatments might be most effective.

**Graphical Abstract:** 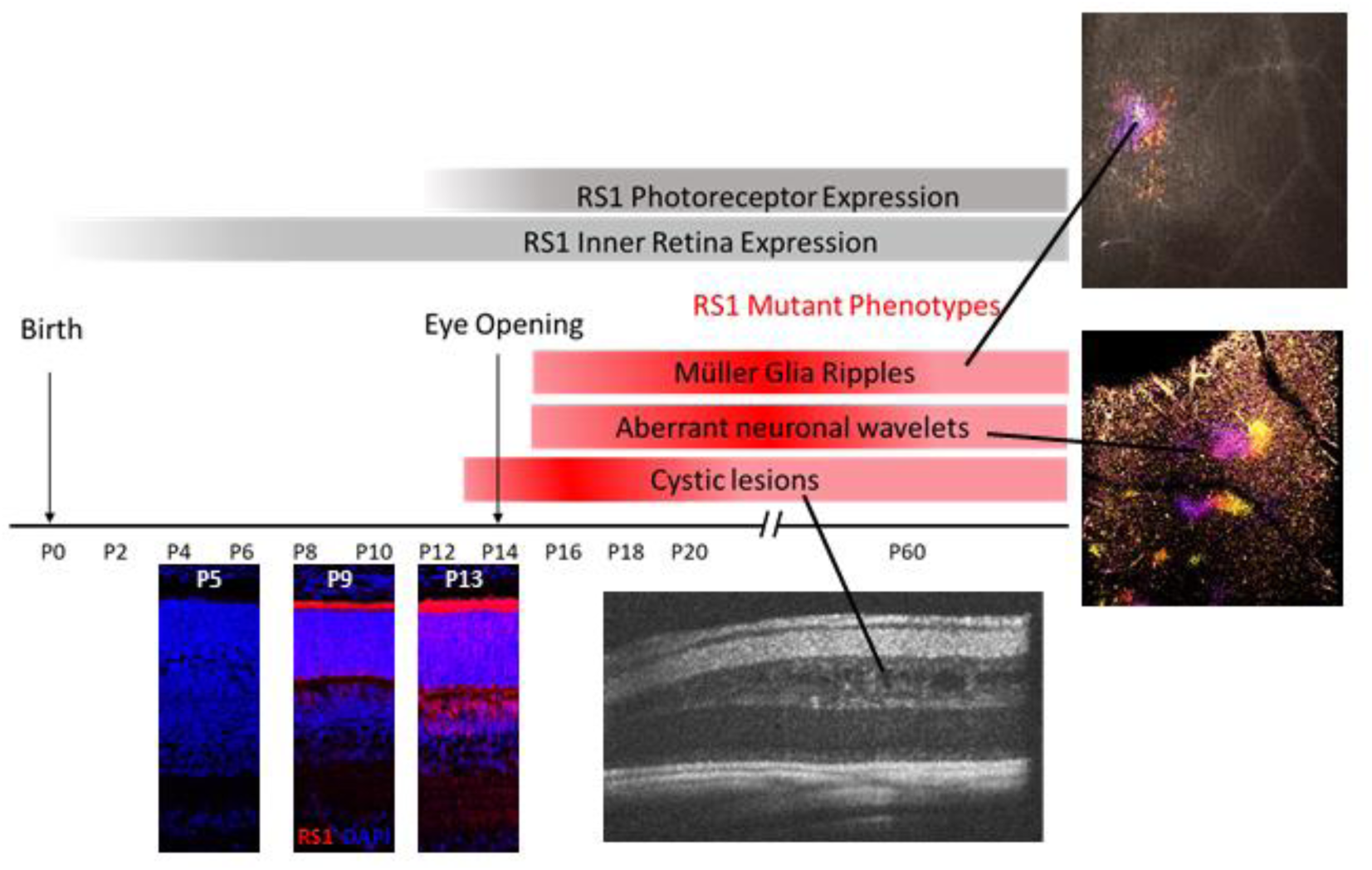

## Introduction

X-linked Retinoschisis (XLRS; OMIM: 312700) is a leading cause of early onset macular degeneration in males. XLRS is characterized by a splitting of the retinal layers (R. S. Molday, Kellner, & Weber, 2012) with cystic-appearing lesions most apparent in the central macula, although approximately 50% of patients also develop peripheral retinoschisis (George, Yates, & Moore, 1995). Functional diagnostics have uncovered significant losses in visual acuity and contrast sensitivity, as well as amplitude reductions of the electroretinogram (Alexander, Barnes, & Fishman, 2005; Forsius et al., 1973; Peachey, Fishman, Derlacki, & Brigell, 1987; Tanino, Katsumi, & Hirose, 1985). XLRS is caused by mutations in the *RS1* gene which encodes retinoschisin (RS1), an extracellular protein exclusive to the retina (Sauer et al., 1997). However, there still exists a number of gaps in knowledge, especially at early developmental stages.

In the mature mouse retina, RS1 is secreted primarily by photoreceptors (Liu et al., 2019). While it is observed in inner retinal cells during development (Liu et al., 2019; Takada et al., 2004), the functional role of this expression is unclear. The protein consists of disulfide bond stabilized homo-dimers, which assembles intracellularly (R. S. Molday et al., 2012) in the form of octamer pairs displaying a symmetrical cog wheel-like structure (Bush, Setiaputra, Yip, & Molday, 2016; Tolun et al., 2016). This rugged topography, along with the anatomical features of retinal splitting in the absence of RS1, is consistent with the proposed role for RS1 as a retinal adhesion molecule (Wu, Wong, Kast, & Molday, 2005). RS1 has further been identified as a binding partner for the α3 β2 isoform of the Na/K ATPase at the surface of photoreceptors and may additionally be regulating intracellular pathways (L. L. Molday, Wu, & Molday, 2007; Plossl, Royer, et al., 2017; Plossl, Weber, & Friedrich, 2017). In recent work (Liu et al., 2019), we noted that horizontal cells (HCs) and rod bipolar cells (RBCs) extended their neurites past the outer plexiform layer (OPL) and into the outer nuclear layer (ONL). These anatomical defects suggest a role for RS1 in controlling glutamate release from photoreceptor terminals. This hypothesis is further supported by functional studies identifying both pre-synaptic and post-synaptic abnormalities at the glutamatergic photoreceptor-to-ON-BC synapse of *Rs1^-/-^* (KO) mice (Ou et al., 2015). The link between defects at the OPL, vision loss and anatomical cyst formation remains to be fully understood.

In our previous work (Liu et al., 2019) we noted severe disease phenotypes in mice aged postnatal day (P) 15 in an allelic series of 3 *Rs1* mutants. In these animals, spontaneous aberrant activity in retinal ganglion cells (RGCs) was observed using patch clamp electrophysiology. While this approach provides information regarding the connectivity of a single neuron, it can only examine the functional properties of healthy cells that can be patched successfully, as it is near impossible to establish and maintain a giga-ohm seal with an intracellular pipette on dystrophic cell membranes. Patch clamp electrophysiology is thus unable to evaluate the dynamics of unhealthy retinal neurons. Multiphoton microscopy of mouse lines expressing the fluorescent calcium indicator GCaMP6f avoids this shortfall and has the additional feature of allowing intracellular calcium dynamics to be acquired simultaneously from cells in all retinal layers. In the present report, we employ calcium imaging optophysiology to better understand the KO phenotype.

Our results revealed aberrant neuronal signals that we termed ‘wavelets’, which persisted after eye opening in KO mice and were found to originate in the OFF-portion of the inner plexiform layer (IPL). Additional aberrant signals were observed in clusters of Müller Cell (MC) glia which displayed spontaneous and coordinated ripples of calcium fluorescence, which propagated radially in both control and KO mice. These glial ripples were significantly larger and more frequent in KO than in control retinas. Abnormal wavelets and glial aberrant signals were driven by glutamatergic neurotransmission. Using *in vivo* spectral-domain optical coherence tomography (SD-OCT) imaging, we report that retinoschisis is clearly present at P13 but was not observed at P11 in three *Rs1* mutant models, indicating that initial retinal development may proceed normally in the absence of wildtype RS1. This time course matched that when RS1 protein was encountered outside of the photoreceptor inner segments.

Our investigations provide a blueprint for the pathophysiological mechanisms associated with mutant RS1, define a key role for neuro-glial cells of the retina, and identify a temporal window for how the onset of dysfunction coincides with RS1 appearance outside of photoreceptor inner segments. By conferring a functional role to RS1, our results strengthen the hypothesis that RS1 is more than an adhesion molecule but is instead essential for the maturation of retinal circuitry. Further, our results suggest an intimate relationship between RS1 and normal glutamatergic synapse formation. Together, these findings identify a set of temporal and cellular targets for novel treatments of XLRS.

## Results

### 1. Developmental time course of *Rs1* expression and RS1 localization

Based on our observations that the disease phenotype is severe in *Rs1* mutants as early as P15 (8), we further characterized the time course for *Rs1* expression and RS1 distribution using *in situ* hybridization and IHC on retinal sections from mice aged P3, P5, P7, P9, P11 and P13. *Rs1* mRNA transcription is first observed at P5, with a steady increase throughout development (**Figure S1A, C, E**). RS1 protein is first clearly detected at P9, in the photoreceptor inner segments and also in the developing OPL, with a subsequent age-dependent increase in the inner segments and throughout the retina (**Figure S1A, B, D**).

### 2. *In vivo* experiments reveal structural and functional abnormalities that initiate at postnatal day 13 for three different Rs1 mutant lines

We examined the developing retina with respect to a key feature of the *Rs1* mutant phenotype, the presence of schitic cavities on OCT imaging. Figure S2 compares representative OCT images of WT and *Rs1* mutant retina at P11 and P13. the time window when RS1 protein is restricted to photoreceptors and then is seen throughout the retina. At P11 (**Figure S2A**), the phenotype of *Rs1* mutant mice is indistinguishable from WT. All retinal layers are present, and there is no evidence of schisis. By P13 (**Figure S2B**), schisis is observed in each *Rs1* mutant model. We previously showed that retinal schisis progresses further to a more severe phenotype at P15 (Liu et al., 2019). In view of the similarity between the phenotypes of the three *Rs1* mutant lines, we further explored this early time window using optophysiological analysis of the GCaMP reporter line crossed to the KO line. We chose the KO as a representative XLRS model to provide reproducible comparison points with existing and future literature as various *Rs1* null mutants have been developed and analyzed by multiple laboratories (Jablonski et al., 2005; Weber et al., 2002; Zeng et al., 2004).

### 3. Generation and validation of a reporter system for the unbiased characterization of neuroglial dynamics

To characterize the intracellular calcium dynamics of retinal cells in the absence and presence of RS1, we generated a mouse line that expresses the fast intracellular calcium indicator GCaMP6f (T. W. Chen et al., 2013) in most retinal cell types (neurons, glia and vascular cells) by crossing two commercially available transgenic lines (**Figure 1A**). The *NG2 Cre* mouse (B6;FVB-Tg(Cspg4-cre) 1Akik/J, #008533, Jackson Labs) expresses *Cre recombinase* under the control of the mouse chondroitin sulfate proteoglycan 4 (Cspg4, also known as Neuro-Glia 2 (NG2)) promoter, resulting in early embryonic *Cre* expression in CNS glial, vascular and neuronal cells (Zhu, Bergles, & Nishiyama, 2008). It should be noted that the NG2 promoter is only active in contractile vascular cells during adulthood (Zhu et al., 2008). The floxed GCaMP6f mouse (B6J.Cg-Gt(ROSA)26Sortm95.1(CAG-GCaMP6f) Hze/MwarJ, #028865, Jackson Labs) has the GCaMP6f construct downstream of the ubiquitous CAG promoter and a stop sequence flanked by loxP sequences, resulting in strong *GCaMP6f* expression by cells also expressing *Cre recombinase*. The resulting offspring (*NG2 GCaMP6f*) express the calcium indicator in all CNS neural, vascular and glial cells (including the retina). We interbred these mice until a fully homozygous line was obtained (*NG2 GCaMP6f)*.

**Figure 1:**
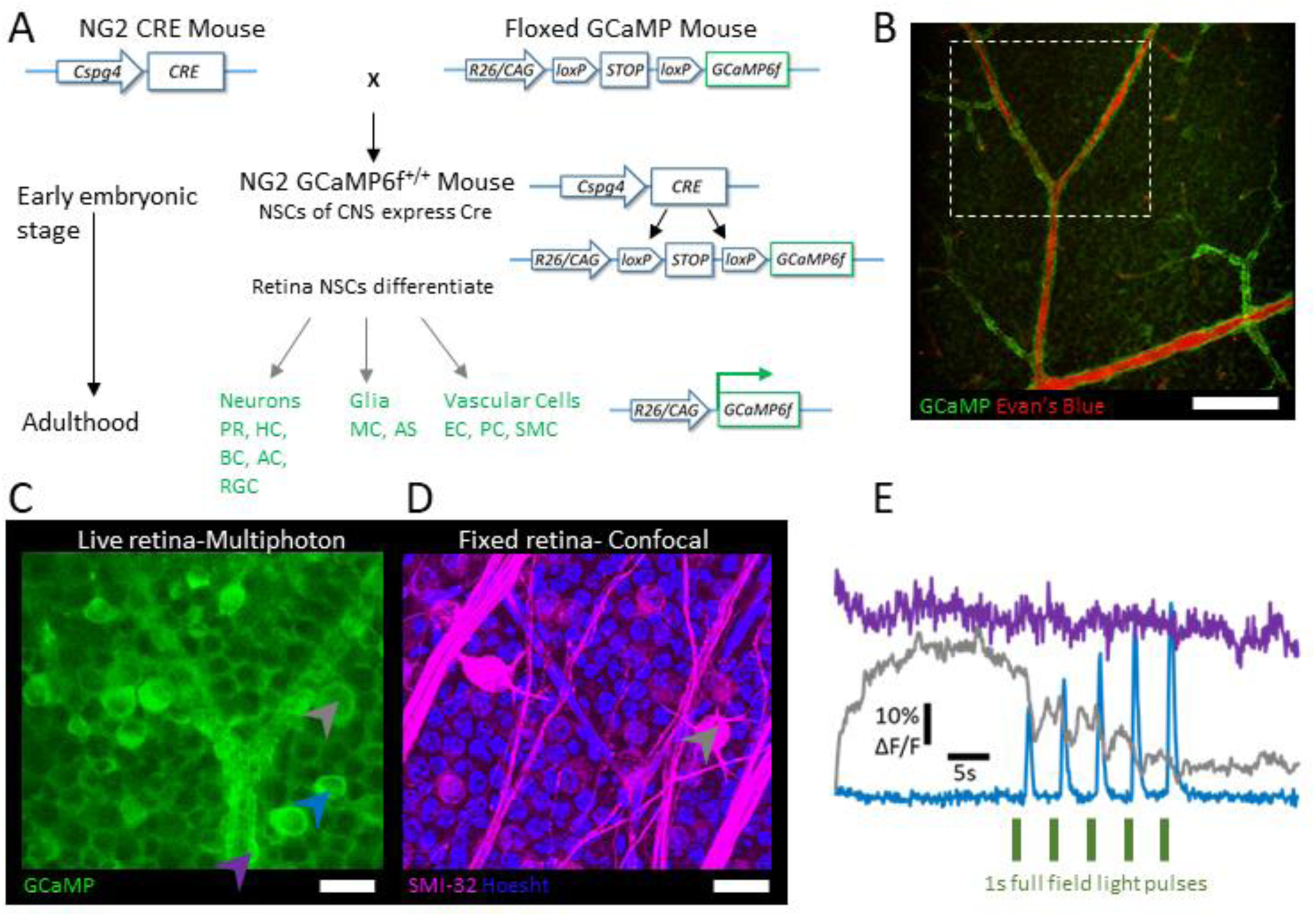
Generating *NG2 GCamp6f* mice to characterize calcium dynamics in retinal cells. **A.** The NG2 Cre parent (left) expresses Cre recombinase under the Cspg4 promoter, active in all neuroglia cells at early embryonic stages. The floxed GCaMP parent (right) expresses a floxed stop cassette upstream of the GCaMP6f sensor construct on the Gt(ROSA)26Sor locus, downstream of the ubiquitous CAG promoter. **B.** Multiphoton imaging of the calcium dynamics in neurons, vascular cells and glia via the green channel simultaneously to mapping the local vascular system (injected with Evan’s Blue) for post-hoc immuno-histochemical analyses via the red channel. Scale Bar is 40 µm. **C.** Portion of retina in (**B**, white box), highlighting several fluorescent retinal ganglion cells (blue and grey arrowheads) and vascular cells (purple arrowhead). Scale Bar is 20 µm. **D.** Retina in **(B & C)** stained for a sub-population of neurons (alpha RGCs, SMI-32, magenta) and nuclei in the GCL. Elongated nuclei of vascular cells indicate the shape of local vasculature. Scale Bar is 25 µm. **E.** 30s fluorescent traces of cells indicated in **(C)** with the neuronal traces (grey and blue) showing fast responses to light stimulation (5x 1s pulses every 5s, green) and the vascular cell (purple) showing no response, but a higher fluorescence baseline. AC: Amacrine Cells, AS Astrocytes, BC: Bipolar Cells, EC: Endothelial Cells, HC, Horizontal Cells, MC: Müller Cells, NSC: Neural Stem Cells, PC: Pericytes, PR: Photoreceptors, RGC: Retinal Ganglion Cells, SMC: Smooth Muscle Cells.

We then crossed *NG2 GCaMP6f* homozygous males with homozygous KO females lacking *Rs1*, resulting in male offspring that were heterozygous for *NG2 GCaMP6f* and lacked RS1 *(NG2 GCaMP6f^+/-^; Rs1^y/-^*). As the *Rs1* gene resides on the X chromosome (Sauer et al., 1997), these male mice, hereafter referred to as KO, exhibited the phenotype observed in *Rs1* mutants including retinal schisis which was visible in our *ex vivo* configuration (Liu et al., 2019). To monitor calcium dynamics in age- and sex-matched control animals, we crossed NG2 GCaMP6f^+/+^ mice with the WT C57BL/6 line.

We used live *ex vivo* multiphoton imaging to record simultaneously from different depths of the retina. Our set-up allowed for concurrent imaging of physiological processes with GCaMP (Green) and three-dimensional navigation of retinal explants via the Evan’s blue labeling of the vascular lumen with Evan’s Blue (**Figure 1B**, red); generating a “fingerprint” of the recording zone, to match with IHC for post-hoc analyses. Individual neuronal cell bodies were readily observable in the retinal nuclear layers, as can be seen in the GCL of a representative retina in **Figure 1C**. Many of these responded to 565 nm light-stimulation with ON, OFF or ON-OFF fluorescence profiles (**Figure 1E,** grey and blue arrowheads). Large alpha-type RGCs were easily recognizable from their appearance (**Figure 1C,** grey arrowhead) and light responses (**Figure 1E,** grey trace), and could be stained by IHC with SMI-32 (**Figure 1D** magenta).

Endothelial cells, pericytes and smooth muscle cells were readily identifiable around blood vessels by their morphology as they displayed strong baseline fluorescence (**Figure 1E,** purple trace). The processes of MCs and astrocytes were observable between neurons and around vascular structures yet were only identifiable when they displayed strong sporadic bursts of fluorescence (characterized below). After fixation, MCs and vascular structures presented a stronger fluorescence signal than did neurons or astrocytes (**Figure S3**).

### 4. Abnormal neural “wavelets” appear at eye opening

We monitored spontaneous and light-driven calcium dynamics using multi-photon calcium imaging in *ex vivo* retinal explants (methods) at ages that correspond to key stages of visual development: P10-11, P14-15, P20-22, and P60-90.

Stage II developmental waves are an important feature of retinal development, refining lateral cholinergic synapses between RGCs and ACs of the GCL (Feller, Wellis, Stellwagen, Werblin, & Shatz, 1996). **Figure 2A, B** display representative Stage II waves at P10, each occurring across an eight second recording epoch and covering over 500 µm. For P10 control mice, the pseudocolor scale shows coordinated rightward propagation in time from 0 (blue) to 7.5 sec (white). To highlight the propagation profile of coordinated waves of activity, the pseudocolor plot is broken down into 3 monochrome snapshots in time within the eight seconds of activity, at time 0, 3s and 6s below the large pseudocolor plot, for each age. In contrast, **Panel B** shows coordinated propagation in the KO mice at all ages. **Panel C** compares normalized fluorescence in control and KO over time, for a representative ROI at each of four ages. Panels **D-G** display summary metrics (Frequency, Distance, Duration and Velocity) for the waves recorded in each explant at these ages. Spontaneous events recorded at P10 exhibited frequency, velocity, duration and inter-wave-intervals typical of Stage II retinal waves (Demas et al., 2006; Maccione et al., 2014). These did not differ significantly between the control and KO mice with respect to any parameter (**Figure 2, D-G, Video 1-2**).

**Figure 2:**
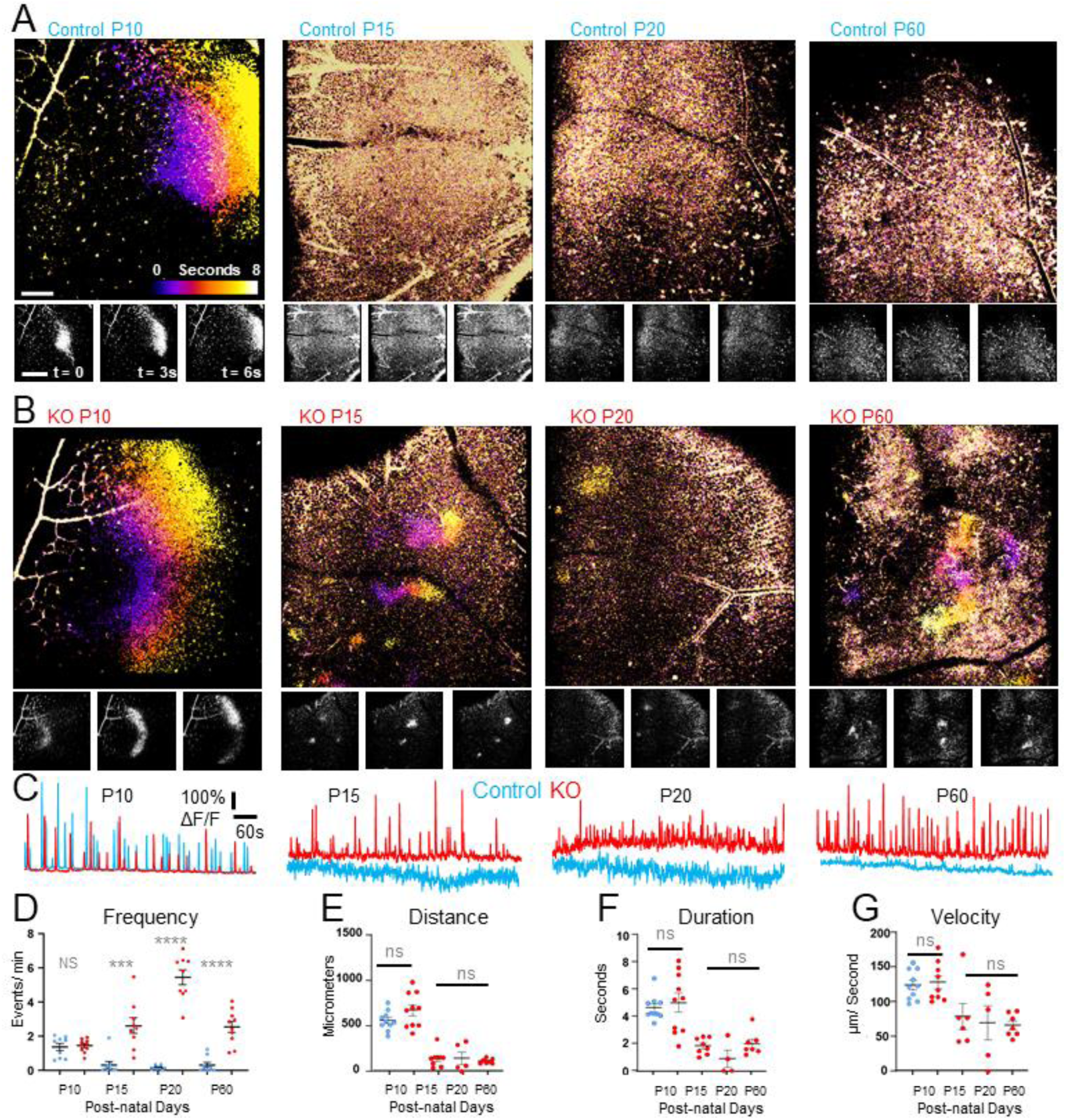
Progressive divergence of spontaneous neuronal activity during retinal maturation. **A.** Representative propagation profiles of P10, P15, P20 and P60 control retinas. While developmental neural waves were observed in controls at P10, they were not seen at later ages. **B.** Representative propagation profiles of P10, P15, P20 and P60 KO retinas. The developmental neural waves seen at P10 persisted at later ages. Scale bar is 100 µm in large temporal projection and 400 µm in small monochrome fames. **C.** Representative fluorescence signal traces from r 100×100 pixels ROIs for each genotype at each age. **D.** Average number of spontaneous events/min/explant, averaged over 5 ROIs. N = 11 (P15 control), 8 (P15 KO), 10 (P20 control), 11 (P20 KO), 8 (P60 control), 12 (P60 KO) from 6 mice per condition. **E-G.** Average distance **(E)**, duration **(F)**, velocity **(G)** for 5 waves/ explant. Mann-Whitney test with multiple comparisons yielded p values of 0.999 (**D**, Frequency), 0.7546 (**E**, Average Distance per explant), 0.616 (**F**, Average Duration per explant), 0.842 (**G**, Average Velocity per explant). For frequency, Mann-Whitney test with multiple comparisons yielded p values of 0.0002 (P15, N = 11 control explants and 15 KO explants), <0.0001 (P20, N = 10 control explants and 11 KO explants), <0.0001 (P60, N = 8 control explants and 12 KO explants). Mann Whitney tests yielded p-values of 0.357 (**E**, Average Distance per explant), 0.877 (**F**, Average Duration per explant), 0.3680 (**G**, Average Velocity per explant).

At later ages, KO retinas exhibited a previously un-observed phenomenon which we termed ‘wavelets’, due to their transient lateral propagation along the neuronal field (**Video 4, Figure 2B-C**). To characterize this novel phenomenon, we applied metrics typically used (Demas et al., 2006; Maccione et al., 2014) for the study of retinal developmental waves (Inter-wave interval distribution, duration, frequency, velocity, distance travelled). We initially looked at large portions of the retina (16x magnification, 734 µm x 834 µm) to ensure that this phenomenon was not restricted to isolated locations, such as cystic cavities. To maximize reproducibility, for each explant we analyzed five different ROIs. Each ROI was 100×100 pixels (122.6 x 122.6 micron) and was chosen pseudo-randomly during post-hoc analysis with the sole criteria that the ROI was devoid of visible vasculature.

After eye opening, we noted that waves stopped in control retinas, as noted previously (Feller, 2009; Feller et al., 1996; Maccione et al., 2014), but persisted as wavelets in KO retinas. **Figure 2A** displays several examples of these occurring over an 8 second period. Although these events were more frequent **(Figure 2, D),** they were shorter, slower and smaller than Stage II waves seen at P10 (**Figure 2, D-E**). Individual wavelet amplitude distributions indicated that the signals recorded from control retinas were negligible compared to those of KO retinas (**Figure S4A**). The average distance, duration and velocity of KO wavelets recorded per explants did not differ significantly between age groups (**Figure 2, E-G**). The individual values for the amplitude, inter-wave interval and durations are plotted as violin distributions (**Figure S4A-C**), demonstrating very similar profiles across age.

### 5. Abnormal Glial phenotype

At P10, control and KO retinas had very similar morphologies, consistent with the similar spontaneous neural dynamics. In comparison, older KO retinas displayed many abnormalities in addition to the cystic structures that develop and characterize the disease phenotype (Liu et al., 2019). Some of those abnormalities were observed as fibrous structures in the plexiform layers (**Figure S5A, C**). These may be due to reactive MCs, which upregulate GFAP in *Rs1* mutant mice (Liu et al., 2019) and many other retinal conditions (Gargini, Terzibasi, Mazzoni, & Strettoi, 2007; Hippert et al., 2015; Yu et al., 2004). In advanced photoreceptor degenerations, MCs are known to drive outer retinal remodeling, displaying hypertrophy, hyperplasia, phagocyting dead photoreceptors and colonizing the sub-retinal space (Jones et al., 2012; Jones, Watt, & Marc, 2005). Using live Z-stack imaging, we characterized the fluorescence-depth profile through a 150 µm retinal column within a pseudo-randomly allocated surface (avoiding vasculature in all layers) of 100 µm^2^ for 6 control (**Figure S5B**) and 6 KO (**Figure S5D**) retinas from distinct animals. In the control retina, the profile has peaks at 20-40 µm and 60-80 µm, possibly corresponding to coordinated activations of the plexiform layers as the descending objective initiated a laser induced contrast response during acquisition. Comparatively, the KO retinas display a stunted distribution, with no isolated peaks, and an overall diminished fluorescence compared to the control retinas.

During the same recordings we analyzed for neural data in **Figure 2**, we observed that clusters of MCs displayed sporadic bursts of spontaneous fluorescence in the adult retina (**Figure 3A, B, Video 5**). These events were identifiable as MC signals because they (1) occurred between neuronal cell bodies, (2) were visible at different depths of the retina (**Videos 7-12**) in a synchronized fashion, and (3) displayed slow and broad temporal dynamics (**Figure 3C, E**) characteristic of glial signals (Rosa et al., 2015). Additionally, this pattern of activity corresponded to the expression pattern of GS (**Figure S3B**), which is only found in MCs in the retina (Bringmann et al., 2009). Of interest, activity was propagated radially to adjacent MCs, in a stereotypical “Ripple” display **(Figure 3B**, **Video 6**, P20 KO displays a particularly sustained example) sometimes propagating over distances of several hundred microns. Hence, we refer to them throughout the rest of this study as “glial ripples”. These glial ripples were rarely seen at P10. In the 20 explants (n = 6 control mice and 6 KO mice, multiple explants used per animals, see methods) investigated at this age, only 3 such glial ripples were observed (1 control, 2 KO). In comparison, every 11 min recording from WT explants at older ages contains 3-5 ripples and those from every KO explant presented more than 5 ripples (Fig. 5D).

**Figure 3:**
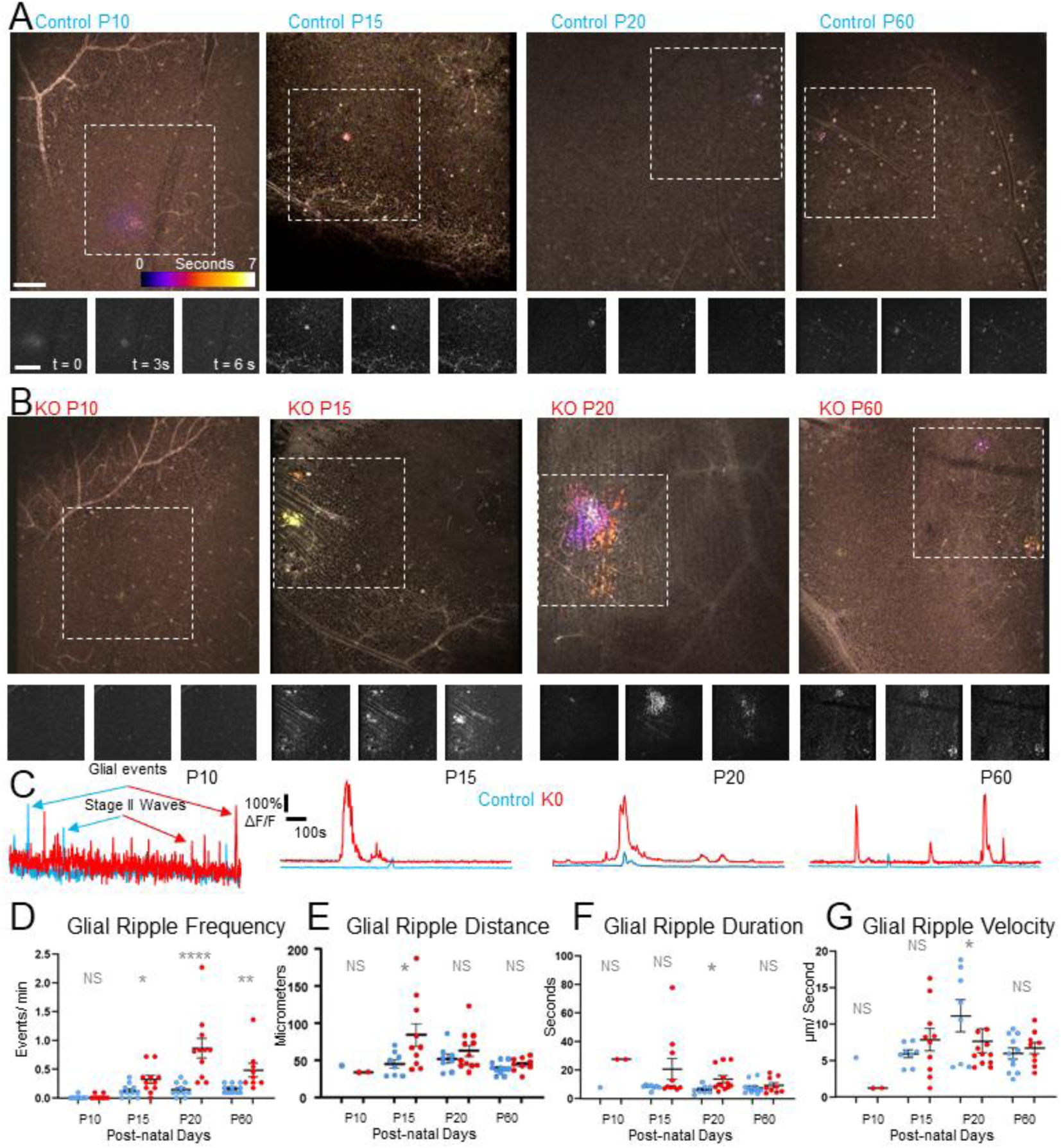
RS1-deficient retinas display aberrant spontaneous glial signals after P15. **A-B.** Spontaneous glial ripple at GCL of control (**A**) and KO (**B**) retinal explants, for each age examined. The three panels at the bottom display still images at key temporal points of the glial ripple. **C.** Example fluorescence traces for both genotypes at key developmental stage. **D.** Frequency of spontaneous glial ripple per explant. **E-G.** Statistical metrics for glial events including velocity **(E)**, duration (**E**) and distance (**F**) for maximum 5 events per explant. N = 11 (P15 control), 8 (P15 KO), 10 (P20 control), 11 (P20 KO), 8 (P60 control), 12 (P60 KO) from 6 mice per condition. **D.** After eye opening, using Mann-Whitney test with multiple comparisons, the p-values at each age when comparing genotypes were 0.028 (P15, N = 11 wt explants and 15 KO explants), <0.0001 (P20, N = 10 control explants and 11 KO explants) and 0.0014 (P60, N = 8 control explants and 12 KO explants). Mann Whitney tests yielded p-values of 0.005 ((**E**, Distance, P15), 0.058 (**E**, Distance, P20), 0.1944 (**E,** Distance, P60), 0.646 (**F**, Duration, P15), 0.0151 (**F**, Duration, P20), 0.710 (**F**, Duration, P60), 0.332 (**G**, Velocity, P15), 0.024 (**G**, Velocity, P20), 0.537 (**G**, Velocity, P60).

Glial ripples were much rarer than the neuronal waves and wavelets characterized in **Figure 2** and appeared at completely different spatial locations. Indeed, ripples almost never occurred in the same location, activating completely different sets of cells each time. As such, we could not rely on an automatic and pseudo-random detection method, so ripples were identified manually (by an observer blinded to genotype and age), and up to 5 ripples per explant were used to calculate average distance, duration and velocity. WT explants displayed frequencies of 0.12 (P15), 0.14 (P20) and 0.16 (P60) ripples per minute, which did not change across age (**Figure 3D**). At each, the ripple frequency observed in KO explants (0.32 (P15), 1.15 (P20), 0.48 (P60) ripples per minute) was significantly higher than in WT, with the highest rate seen at P20 (**Figure 3D**). Other measures of glial ripples (duration, velocity, distance) were more variable and did not always reach statistical significance (**Figure 3D, E, F**). KO ripples tended to propagate over further distances than age-matched controls (**Figure 3E**), and also tended to last for longer durations (**Figure 3F**).

### 6. Aberrant neuro-glial dynamics are driven by dysfunctional glutamatergic signaling

We next sought to characterize the functional and chemical origin of the aberrant neuronal wavelets observed in KO retinas. To locate their origin, we performed four-dimensional simultaneous (FastZ, *cf* methods) imaging of all retinal layers. We used a simple full-field light stimulus (5 pulses, 1s ON, 4s OFF) to identify the ON and OFF portions of the IPL. Spontaneous (non-light- or drug-driven) recordings 9 minutes long allowed us to image the activity in 6 different layers at an effective sampling frequency of 1.975 frames per second. Representative examples of such recordings for both KO and control P60 retinas under standard (vehicle) perfusion conditions are shown in the left portions of **Figure 4A-B**. In control explants taken at P60, very little wavelet activity was seen in the INL, IPL or GCL (**Figure 4A**). Consistent with prior results, KO wavelet activity was elevated, especially throughout the IPL and GCL, with less difference seen in the INL (**Figure 4B, C**).

**Figure 4:**
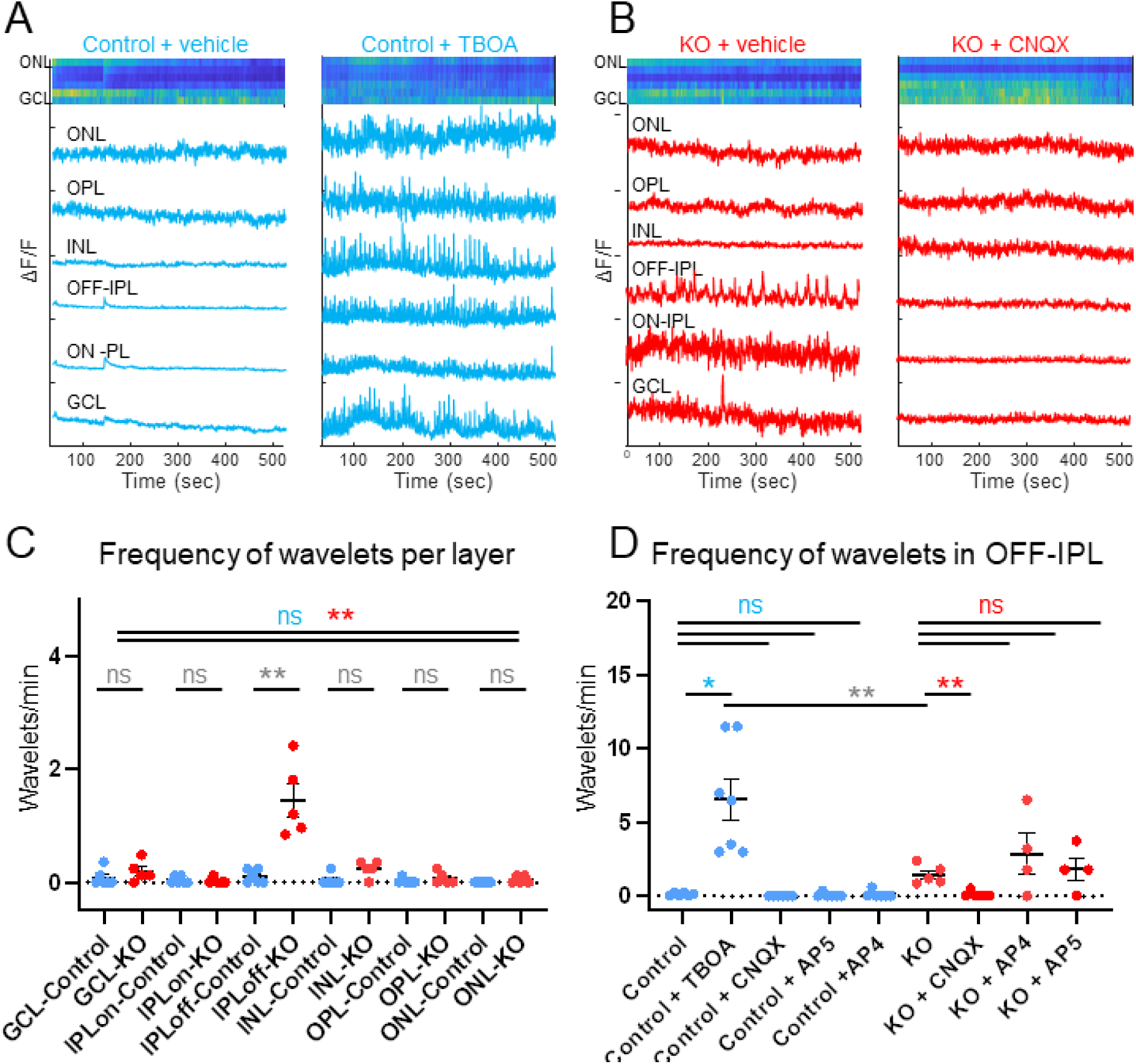
Neural wavelets originate in the OFF-IPL and are driven by glutamate in the mature retina. **A-C.** Example traces from 9 minute recordings of spontaneous activity using FastZ imaging in 6 retinal layers simultaneously in mature control (**A**) and KO (**B**) explants. The ionotropic glutamate receptor blocker CNQX abolished spontaneous wavelets in the off IPL of KO retinal explants (**B**) but the glutamate transport blocker TBOA induced spontaneous signals in control retinas (**A**). **C.** Wavelet frequency was highest in the OFF-IPL layer of KO retinas, and completely absent from control retinas. N = 5 for each genotype. Mann Whitney test, N = 10 animals, 5 per genotype, p = 0.0025 for IPL OFF. **(D)** Wavelet frequency in the off IPL of CONTROL retinas was increased by application of TBOA. Wavelet frequency in the off IPL of KO retinas was reduced by application of CNQX, but not AP4 or AP5. Friedman Test with repeated measures, N = 7 (CONTROL + TBOA, p = 0.0458), N = 5 (CONTROL + CNQX, p =0.0005 & RS + CNQX, p = 0.0043), N = 4 (CONTROL + AP4, p = 0.0041; KO + AP4, p = 0.5556), N = 4 (control + AP5, p = 0.0036, KO + AP5, p = 8571).

Glutamate is the main excitatory neurotransmitter in the retina, with very clear anatomical and functional distinctions in receptor subtypes. We studied the P60 retina using well-defined pharmacological agents to block each of the main glutamate receptor subtypes. CNQX (6-cyano-7- nitroquinoxaline-2,3-dione) is a competitive AMPA/kainate receptor antagonist capable of blocking all ionotropic glutamate receptors, including those on OFF bipolar cells (Diamond & Copenhagen, 1993), which are responsible for the propagation of OFF responses from the OPL to the OFF-portion of the IPL. L-AP4 (L-2-amino-4-phosphonobutyric acid) is a selective agonist for group III metabotropic glutamate receptors, and used in the retina for blocking mGluR6 receptors on ON-bipolar cells, which are responsible for the propagation of ON responses from the OPL to the ON-portion of the IPL (Nakajima et al., 1993). D-AP5 ((2R)-amino-5-phosphonovaleric acid) is a selective antagonist for NMDA glutamate receptors, expressed in the inner retina, on RGCs and ACs (Kalloniatis, Sun, Foster, Haverkamp, & Wassle, 2004). Only CNQX significantly eradicated the aberrant wavelets (**Figure 4D**). The frequency of recorded neural wavelets in the OFF portion of the IPL was significantly smaller in the presence of 10 µM CNQX (Mann Whitney test, p = 0.0043, N = 5 animals). While AP5 (50 µM) and AP4 (25 µM) had potentiating effects on the frequencies of recorded waves, the changes were more variable and not statistically significant (Mann Whitney test, p = 0.5556 for L-AP4 and p = 0.8571 for D-AP5, N = 8 animals). We also used TBOA (DL-threo-beta-benzyloxyaspartate), to block excitatory amino acid transporters (EAAT1) located on MCs (Derouiche & Rauen, 1995) which are responsible for the first step in the recycling cascade of glutamate. While TBOA induced wavelet-like events, these were present in all retinal layers (**Figure 4B**), thus greatly exceeding the pattern seen in the KO retina.

Unlike neuronal wavelets, glial ripples propagated radially throughout all layers of the retina (**Figure 5A**) with the peak fluorescence intensity for each layer phase-locked throughout the retina (**Figure 5D**). This trans-retinal coordination is consistent with a MC origin of the signals. Throughout the retina, the frequency of glial ripples in KO retinas was significantly higher than in control retinas under standard (vehicle) perfusion with aCSF (**Figure 5B**). The addition of the glutamate receptor blockers CNQX, AP5 and AP4 to the perfusion did not alter the frequency of glial ripples in either control or KO retinas. In control retinas, perfusion of GLAST blocker TBOA did not alter the frequency of ripples throughout the retinal layers. Surprisingly, glutamatergic blockers significantly altered the surface area covered by glial ripples in both control and KO retinas, with mixed effects depending on the blocked glutamate receptor. This was also the case for TBOA, confirming a link between glutamatergic signaling and the kinetics of glial ripples (**Figure 5C**).

**Figure 5:**
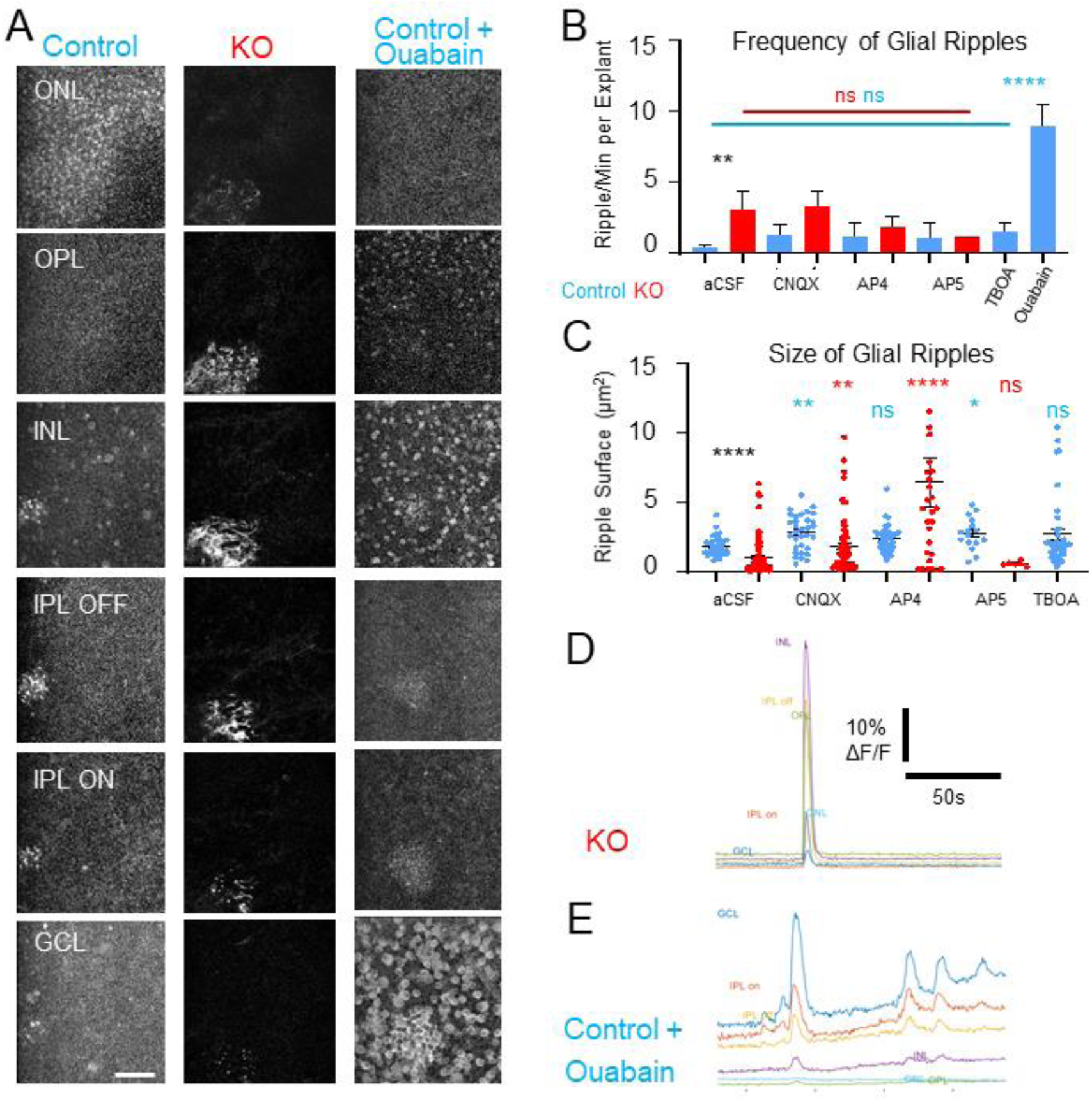
Spontaneous glial events occurred throughout the retina and can be modulated pharmacologically in the mature retina. **A.** Example spontaneous (Control, KO) and induced (Ouabain) glial events within all layers of the retina (P60), projected over 40s. **B**. Throughout retinal layers, the frequency of glial ripples in KO retinas was significantly higher than in control retinas (Mann-Whitney test, p = 0.0052, N = 16 control and 11 KO explant, aCSF KO vs Control). Glutamate blockers did not modulate the frequency of glial ripples in either control (Kruskall Wallis test, p = 0.1830, N = 6 explants per retina per condition) or KO retinas (Kruskall Wallis, p = 0.8989, N = 5 explants per each of retina per condition). Ouabain was a potent inducer of glial ripples (Mann-Whitney test, p <0.0001, N = 5 explants for each of 5 retinas per condition, Control Ouabain vs aCSF). **C.** Throughout retinal layers, the size of glial ripples in KO retinas was significantly smaller than in control retinas (Mann-Whitney test, p <0.0001, N = 16 explants for 6 control retinas and 11 explants for 5 KO retinas). Glutamatergic blockers significantly altered the surface area covered by glial ripples in both control and KO retinas (Kruskall Wallis with multiple comparisons against aCSF: for CNQX, p = 0.0028; for AP4, p = 0.0595; for AP5 p = 0.0216 ; for TBOA, p >0.9999 , N = 5 explants for each of 5 mice per condition). **D-E**. Example fluorescent traces throughout the layers for ROIs displayed in **(A)** for a representative KO **(E)** and a control **(D)** explant treated with ouabain.

We were able to induce high frequency ripples throughout retinal layers in the control retina by applying ouabain, an antagonist for the Na^+^/K^+^ ATPase. Increased intracellular sodium reduces the activity of the sodium-calcium exchanger (NCX), which pumps one calcium ion out of the cell and three sodium ions into the cell down their concentration gradient. This leads to an increase in intracellular calcium, and eventual death of the cell/tissue, in a very repeatable and sequential manner, ending with high-frequency glial ripples (**Figure 5B, D-E, Videos 13-18**).

## Discussion

### 1. Schisis, Neuronal Wavelets, Glial Ripples, Glutamate-dependency

Following our prior observation that *Rs1* expression is initiated between postnatal days 7 and 14 (Liu et al., 2019), we sought to investigate markers of dysfunction associated with the lack of the RS1 protein at these early stages. We first confirmed *Rs1* expression and detectable RS1 protein in photoreceptor inner segments at P5 and P9, respectively. We then identified both structural and physiological phenotypes in *Rs1* mutant mice from postnatal day 13, indicating a four-day lag in the role for the protein in retinal function and structure. These phenotypes presented in the forms of cystic lesions observable by OCT *in vivo*, and the appearance of aberrant wavelets of neural activity. Interestingly, all three *Rs1* mutant mouse models (KO; C59S; R141C) displayed a similar temporal appearance of schisis, consistent with these specific point mutations as well as null mutations being linked to human disease ((R. S. Molday et al., 2012) – review of human mutations). Our further investigations into the mechanisms underlying the novel phenotype of aberrant wavelets in the KO revealed a link to glutamate signaling, a key neurotransmitter in retinal development and function.

### 2. Calcium imaging for dissecting retinal function and dysfunction

Most functional studies of *Rs1* mutant mouse lines have focused on whole animal measures (OCT imaging, electroretinography) that provide links to key phenotypic features of XLRS patients (D. Chen et al., 2017; Jablonski et al., 2005; Weber et al., 2002; Zeng et al., 2004). We previously reported the first evidence of aberrant spontaneous activity in RGCs using single-cell patch clamp electrophysiology (Liu et al., 2019). The present comprehensive optophysiological dataset allows us to investigate the intricacies of physiological activity upstream of these hyperactive RGCs, and concurrently in neuro-glial cell populations. Imaging of intracellular calcium dynamics has been essential for exposing and dissecting spontaneous developmental waves (Meister, Wong, Baylor, & Shatz, 1991). The advent of multiphoton imaging has expanded our ability to characterize RGC and bipolar cell function (Baden et al., 2016; Baden, Euler, & Berens, 2020; Franke et al., 2017). In parallel, an explosion in the availability of selective promoters and transgenic mouse lines has enabled the targeting of distinct cell populations. Recent examples include the imaging of retinal MCs during development (Rosa et al., 2015) and vascular signaling in the retina , leading to a 3D mapping of the pericyte connectome (Alarcon-Martinez et al., 2020; Kovacs-Oller, Ivanova, Bianchimano, & Sagdullaev, 2020). To better understand the KO phenotype we developed a new *Rs1*-deficient mouse line with ubiquitous expression of a fast intracellular-calcium signaling sensor. Of note, we documented the coordinated activation profile of connected columns of MCs in both the healthy control and KO retinas (glial ripples). The further characterization of these events and their underlying physiological relevance in these and other models should help elucidate some of the mysteries underlying functional hyperemia by the neuro-glial-vascular unit. Additionally, we identified spontaneous bursting in vascular cells during development, that was inversely correlated to spontaneous developmental GCL waves (**Video 1, Figure 6**). We are currently working to develop this novel and intriguing line of inquiry.

**Figure 6:**
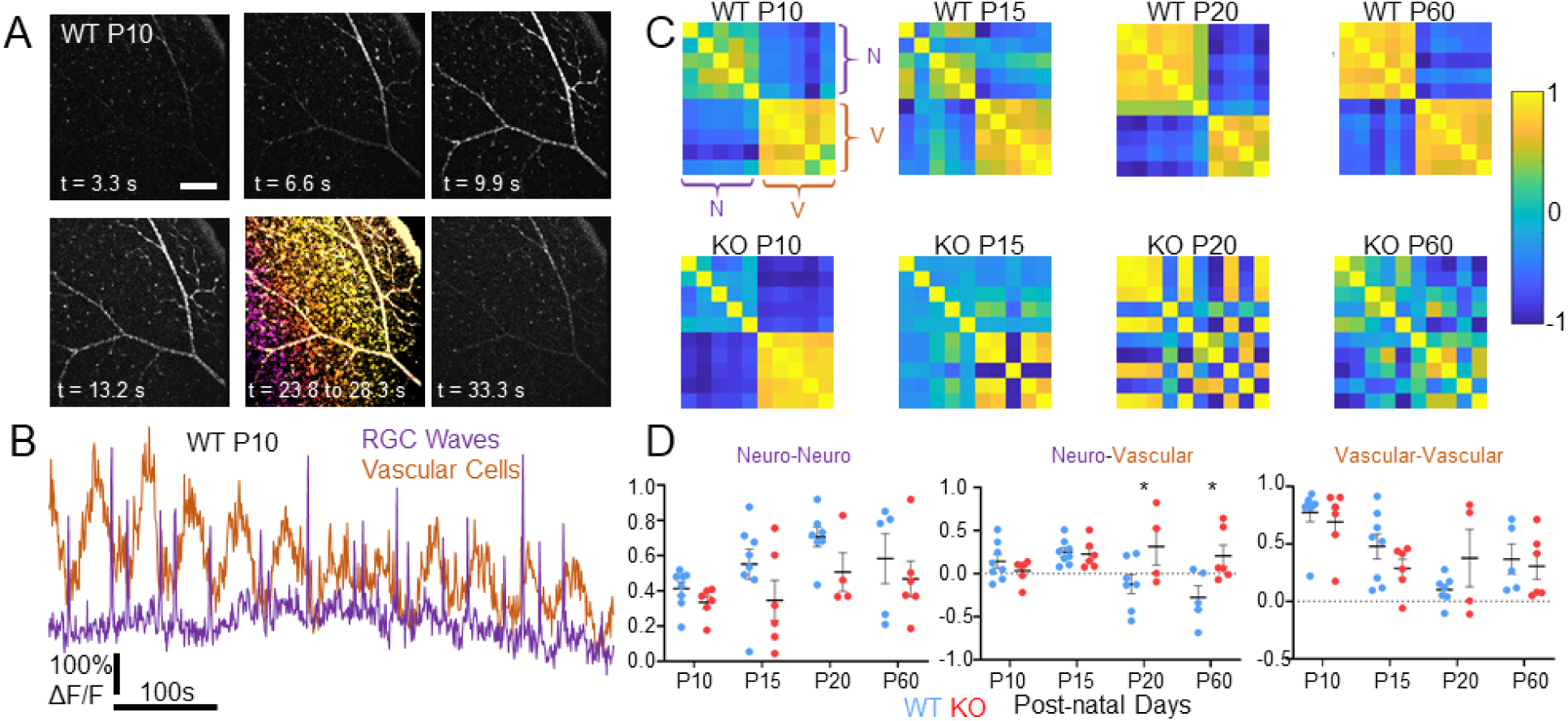
Spontaneous activity in in vascular cells is anti-correlated to spontaneous waves of neuronal activity at P10. **A.** Example spontaneous wave of neuronal activity in the ganglion cell layer intercalated by waves of vascular activity. Scale Bar is 200 µm. **B.** Trace of vascular and neural signal in adjacent regions of interests in (**A**)**. C.** Example correlation coefficient matrices of 5 vascular and 5 neuronal regions of interest for WT and KO at key stages of visual development. For WT and at P10, the vascular (v) activity is highly auto-correlated between regions. **D.** Average correlation coefficients from 5 ROIs in KO and WT retinas at key developmental ages (WT Ns = 8, 8, 7, 5; KO Ns =6, 6, 4, 6 for P10, P15, P20 and P60, respectively). Neuro-vascular correlations are significantly different between WT and KO at P20 and P60 (2-way ANOVA with Sidak’s multiple comparisons, P = 0.032 for P20 and P = 0.011 for P60).

### 3. Role of RS1 in development

It is well established that *Rs1* is almost exclusively expressed in photoreceptors and that the RS1 protein is localized throughout the retina (Takada et al., 2004). Here we examined the developmental time course over which these features develop, towards clarifying the cellular events that accompany the earliest functional and structural defects. Our dataset supports the premise that RS1 is more than just an extracellular adhesion protein maintaining retinal integrity. Previous studies identified the β2 subunit of the retina-specific Na/K-ATPase as an interacting partner for retinoschisin (L. L. Molday et al., 2007), essential for anchoring it to the plasma membranes (Friedrich et al., 2011) of photoreceptors. More recent work has demonstrated that RS1 acted as an anti-apoptotic regulator of mitogen-activated protein kinase (MAPK) signaling in the retina (Plossl, Weber, et al., 2017), via its interactions with Na/K-ATPase (Plossl, Royer, et al., 2017). Our data provide a multi-cellular and developmental context to these molecular findings. Specifically, that RS1 plays a key role in the highly coordinated appearance of distinct anatomical and physiological phenotypes with the maturation of glutamatergic synapses at the IPL and OPL (**Figure 8**). Further, they identify an early age range in which RS1 may have additional important protein interactions.

**Figure 7:**
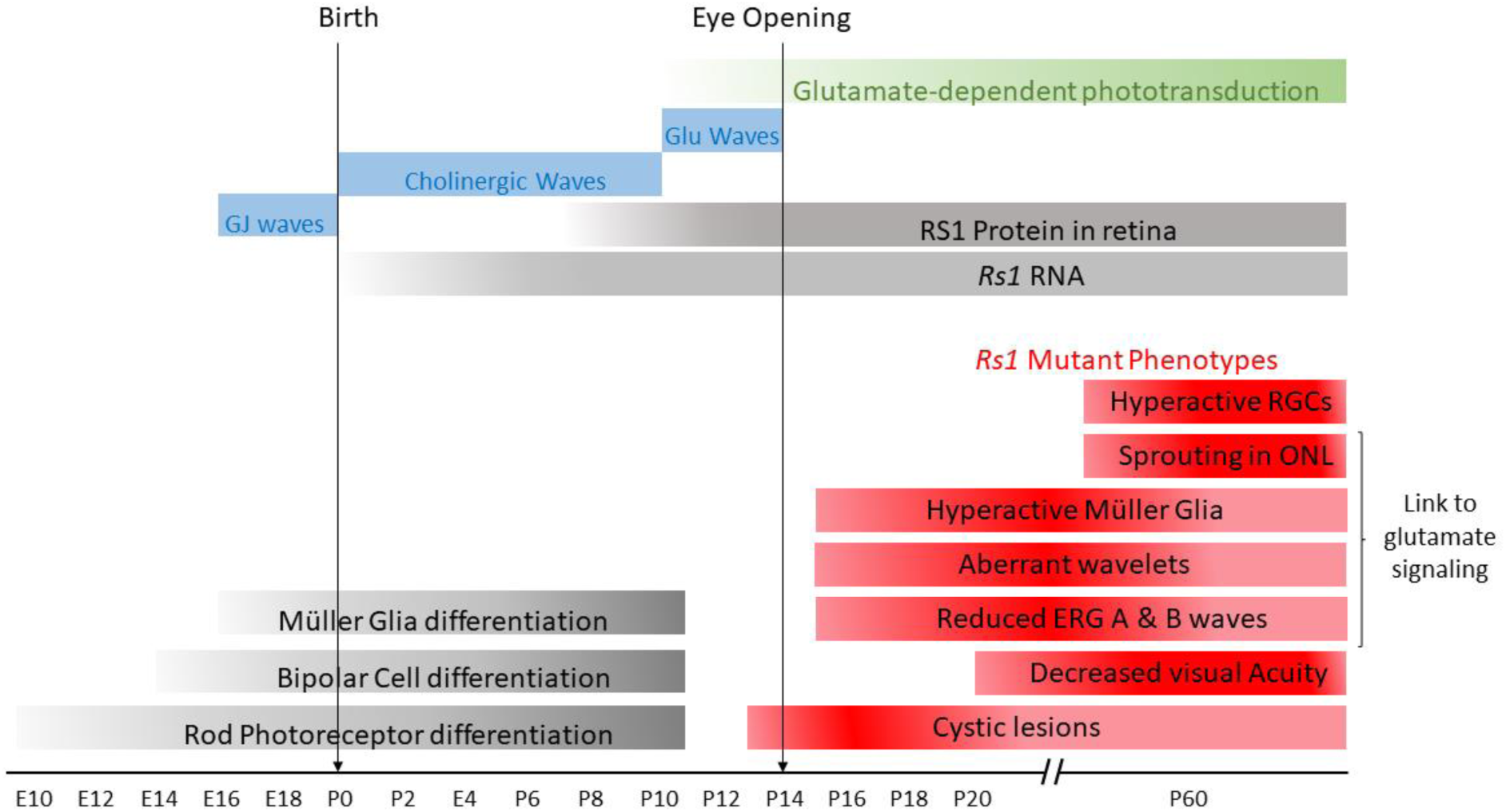
Lack of RS1 protein drives excess glutamate and aberrant wavelets. Coincidence of mutant *Rs1* phenotypes with maturation of glutamatergic synapses. Some of the phenotypes observed in models of XLRS have been linked to glutamatergic signaling. These bars are all based on available data.

In mice, P11 marks the end of rod, BC and MC differentiation (Young, 1985) as well as the onset of the maturation stage for the photoreceptor-to-BC synapses (Stage III waves) which flood the retina with glutamate (Bansal et al., 2000) and prime the visual system for visual perception at eye opening (P15). Our data suggest that this process is disrupted in the absence of RS1, building on previous work that highlighted anatomical (Liu et al., 2019) and physiological (Ou et al., 2015) disruptions at the OPL. For example, Ou et al. (2015) found pre- and post-synaptic defects at the photoreceptor-to-BC synapse in mice models of XLRS. And our recent study noted ectopic neurites from bipolar and horizontal cells (Liu et al., 2019).

### 4. Implications for Development and Repair Strategies

For future pre-clinical studies, our findings provide a robust functional assay independent of macular cysts, whose temporary reversal with the use of carbonic anhydrase inhibitors in XLRS patients has not yielded sustainable clinical improvements in perception. (Ambrosio et al., 2021; Pennesi et al., 2018; Testa et al., 2019; Thobani & Fishman, 2011). For example, drug discovery efforts may focus on the normalization of wavelets or glial ripples without affecting retinal processing in the mature *GCaMP^+/-^; Rs1^y/-^* retina.

Disruption of the spontaneous waves that guide normal visual development leads to a disruption of RGC projections to central targets. Indeed, waves are essential in the wiring of RGC receptive fields, begetting the maturation of their (Stage I) electrical synapses, (Stage II) cholinergic synapses and (Stage III) glutamatergic synapses (Feller, 2009; Feller et al., 1996; Maccione et al., 2014). Our findings that glutamatergic transmission is disrupted in concordance with stage III waves strongly suggest a potential for desegregated inputs to central targets and potentially irreversible connectivity issues. Gene replacement therapy has shown encouraging normalization of outer retinal function and structure in *Rs1* KO mouse models (Dalkara et al., 2013; L. L. Molday et al., 2006; Ou et al., 2015; Takada et al., 2008), leading to a retinal AAV8-RS1 gene therapy phase I/IIa clinical trial (Cukras et al., 2018). Unfortunately, this clinical trial appears to have failed to replicate the observations made in mouse models and has additionally documented that the therapeutic vector induces a considerable inflammatory response (Mishra et al., 2021). With an age range of 23 to 72 years across 9 participants, the results of this clinical trial may instead highlight the need for RS1 in development to generate fully functional visual systems. Future work in whole animal studies might include characterization of the role played by RS1 in support of RGC mapping to central targets (Demas et al., 2006). Investigations in human patients’ central visual pathways may shed some light onto this issue, and identify image forming disparities due to aberrant wiring.

## Methods

### Animals

During all experimental procedures, animals were treated in compliance with protocols approved by the Institutional Animal Care and Use Committees (IACUCs) of Weill Cornell Medicine (WCM), Cleveland Clinic (CC), or Regeneron Pharmaceuticals (RP). All animal studies were conducted in accordance with the National Institutes of Health *Guide for the Care and Use of Laboratory Animals*. For *in vivo* imaging conducted at CC, we examined three *Rs1* mutant lines: a KO with inserted lacZ reporter gene (*Rs1^-/Y^; Rs^-/-^*), a C59S point mutant substitution (*Rs1^C59S/Y^; Rs1^C59S/C59S^*; hereafter C59S), and an R141C point mutant substitution (*Rs1^R141C/Y^; Rs1^R141C/R141C^*; hereafter R141C) (Liu et al., 2019). Controls were an F1 hybrid (F1 hybrid 129S6SvEvTac/C57BL6NTac). For the developmental analysis of *Rs1* expression and RS1 localization conducted at RP, we worked with KO and C57BL/6 mice obtained from Jackson Laboratories and Taconic Biosciences. For optophysiology and immunohistochemistry (IHC) studies conducted at WCM, we crossed KO females with males from our NG2-GCaMP6f line generated by crossing a floxed GCaMP6f line with NG2 Cre transgenic mice (Kovacs-Oller et al., 2020). Control animals were a cross between C57BL/6J females and male NG2 GCaMP6f. Only male offspring (either KO or wildtype) who carried the NG2 Cre and floxed GCaMP6f alleles were used for optophysiological experiments.

### RS1 expression, tissue collection

Following sacrifice, a corneal burn at the superior 12 o’clock position was made to mark orientation. Enucleated eyes were fixed in 4% PFA for 24 hours at 4°C, and a cut was made in the central cornea to allow the fixative to enter the posterior chamber. Following fixation, eyes were washed three times for 15 minutes in 1X PBS. The anterior chamber (cornea and iris) of each eye was dissected and removed, and a small cut was made superiorly at the corneal burn to keep track of orientation. Posterior eyecups were then incubated in 15% sucrose, then 30% sucrose solutions, each overnight at 4°C. Cryoprotected eyecups were embedded in a cryomold containing a 2:1 solution of Tissue-Tek O.C.T. compound (Sakura Finetek) and 30% sucrose, and flash frozen in 2-methylbutane cooled by dry ice. During embedding, the orientation of the eyecup in the mold was kept consistent, with the posterior pole at the bottom of the mold and the optic nerve facing away from the label. Frozen tissue blocks were stored at -80°C until cryosectioning. Ten µm thick serial sections at the level of the optic nerve were collected with a cryostat (Leica CM3050S) onto charged slides (Superfrost Plus Micro Slides, Cat: 48311-703, VWR). The tissue block was oriented so that sagittal sections through the nasal-temporal plane were collected. Collected sections were allowed to dry for 1 hour at room temperature, and then stored at -80°C until used.

### RS1 expression, immunohistochemistry

To prevent non-specific signal from blood vessels by the anti-mouse secondary antibody, 100 µg of the primary antibody against RS1 protein (Abnova, Cat: H00006247-B01P) was directly conjugated to Alexa Fluor 647 (hereafter RS1-AF647) using a site-specific antibody labelling kit (SiteClick™ Antibody Azido Modification Kit, Cat: S0026; SiteClick™ Alexa Fluor™ 647 sDIBO Alkyne, Cat: S10906, ThermoFisher Scientific). Following the final wash step in the protein concentrator, the final volume was concentrated to 100 µl, to achieve an approximate antibody concentration of 1 µg/µl. Sections were blocked in a blocking buffer containing 5% goat serum (Cat: 16210-064, Gibco), 1% bovine serum albumin (Cat: A4503, Sigma) and 0.5% Triton-X (Cat: 93443, Sigma) for 1 hour at room temperature. Sections were then incubated with the RS1-AF647 antibody in blocking buffer (1:100 dilution) overnight at 4 °C. Following incubation, slides were washed three times for 5 minutes with 1X PBS containing 0.1% Tween- 20 (Cat: P9416, Sigma), and two times for 5 minutes with 1X PBS. Excess PBS was carefully removed, and sections were mounted with an anti-fade mounting medium with DAPI counterstain (ProLong™ Gold antifade mountant with DAPI, Cat: P36931, ThermoFisher Scientific) with no. 1.5 glass coverslips (Cat: 106004-340, VWR).

### RS1 expression, in situ hybridization

Slides were washed in distilled water, dipped in 100% ethanol, and allowed to dry overnight at room temperature. Sections were then pretreated with H_2_O_2_ (Cat: 322330, A.C.D) for 10 minutes at room temperature, boiled in antigen retrieval buffer (10X diluted in dH2O, Cat: 322001, A.C.D) for 15 minutes in a steamer (Oster Steamer Model 5709, IHC World), and treated with Protease Plus (Cat: 322330, A.C.D) for 30 minutes at room temperature. At this stage, an additional DNAse treatment step was included to reduce potential background from the probe binding to chromosomal DNA. A solution of DNAse I (50u/ml in 1X DNAse buffer, AM224, Ambion) was added to the sections and incubated at 40°C for 30 minutes. Sections were washed in distilled water 3 times and hybridized with an RNAScope® probe recognize mouse RS1 mRNA (Cat: 456821, A.C.D). The remainder of the protocol was implemented according to the manufacturer’s protocol (RNAScope® 2.5 HD Detection Reagent RED, Cat: 322360, Advanced Cell Diagnostics). Following the final wash step, slides were counterstained with DAPI (1:250 in 1X PBS, 10 mins at RT, Cat: 564907, BD Pharmingen) and mounted with Prolong™ Glass anti- fade mounting medium (Cat: P36980, ThermoFisher Scientific) using no. 1.5 glass coverslips (Cat: 106004-340, VWR). Slides were allowed to dry at room temperature overnight prior to imaging.

### RS1 expression, imaging

Confocal images were acquired with a Zeiss LSM710 line scanning confocal microscope using a Zeiss Plan Apo 20X/0.8 objection and Multi-Alkali Photomultipliers, acquired at 16 bit with a spatial resolution of 3340x3440 pixels. Quantification of RS1 protein and mRNA signal was obtained using FiJi (NIH) following importation of native .czi files. For RS1 mRNA sections, which were acquired at multiple z-planes, an average intensity z-projection was applied to produce a single image from which signal was quantified. In immunohistochemistry images, signal was quantified directly from the raw, single z-plane image.

Regions of interest (ROIs) were drawn at various retinal locations: ganglion cell layer (GCL; ∼7660 µm^2^), inner plexiform layer (IPL; ∼960 µm^2^), inner nuclear layer (INL; ∼7,223 µm^2^), outer plexiform layer (IPL; ∼114 µm2), outer nuclear layer (ONL; 7480 µm^2^), and inner/outer segments (IS/OS, 113 µm^2^). At earlier timepoints (P3 and P5), there is a single retinal layer termed the neuroblastic layer (NBL), and outer and inner nuclear layers have not yet formed two distinct cell layers within the retina. However, even at these early timepoints, cells fated for the inner nuclear layer seem to localize to the anterior (closer to the vitreous) portion of the NBL (Takada et al., 2004). As such, at these early timepoints, the INL was sampled as the anterior portion of the NBL, while the ONL was sampled as the most posterior portion of the NBL. The OPL does not form until P7. For immunohistochemistry images, the OPL was sampled from P7 onwards, and for RS1 mRNA images, from P9 onwards. Mean pixel grey values within the ROIs were obtained and recorded. An additional large ROI (7,024 µm^2^) was drawn spanning the entire retina and a plot of distance versus mean grey value was recorded to obtain a profile of RS1 protein or mRNA expression throughout the entire retina.

### Spectral-domain optical coherence tomography

Imaging procedures have been described in detail (Bell et al., 2015; Bell, Kaul, & Hollyfield, 2014). Mice were anesthetized (sodium pentobarbital, 68 mg/kg) and pupils dilated with 1 µl eyedrops comprised of 0.5% tropicamide and 0.5% phenylephrine HCl. The corneal surface was anesthetized with a single application of ∼10 µl of 0.5% proparacaine. SD-OCT (Envisu R2210 UHR Leica Microsystems Inc.). images of the retina were collected along the horizontal and vertical meridians centered on the optic disk. Each volumetric scan consisted of five B-scans (1000 A-scans per B-scan by 30 frames) which were co-registered and averaged using InVvoVue software (Leica Microsystems Inc.). Each B-scan was ∼1.8 mm in width and had an axial resolution of 1.4 µm.

### Optophysiology, tissue preparation

20 minutes prior to euthanasia by C0_2_, mice were injected intra-peritoneally with a solution of Evans Blue (Sigma Aldrich); 50 µl (33 µg/µl) for animals aged P10-22 and 150 µl (66 µg/µl) for ages >P60. This compound adheres to the vascular lumen and fluoresces in response to a broad excitation spectrum with red-shifted emissions, allowing us to image blood vessels in a different channel to the green-emitting calcium dynamics of GCaMP. For all experiments, dissections were performed in a dark room and the tissue submerged in oxygenized HEPES-buffered extracellular Ringer’s solution, containing the following (in mM): NaCl (137), KCl (2.5), CaCl_2_ (2.5), MgCl_2_ (1.0), Na-HEPES (10), glucose (28), pH 7.4. During imaging, retinal explants were superfused with carboxygenated Ames solution (Sigma Aldrich).

Following enucleation, left eyeballs were immersed in buffered HEPES, pierced with a 3/8 inch 26 gauge intradermal needle (Precision Guide, USA) at the interface between sclera and cornea, then sheared with Vannas scissors along this interface, just below the *ora serrata*. Retinas were teased from the eyecups by pulling on the vitreous with Dumont #5 forceps, then sheared in three or four equitable “slices”, with Vannas scissors. The remaining vitreous humour was removed from the surface of the retina without ever touching the neural tissue. Individual explants were mounted GCL-up onto filter holders (as described previously (Ivanova et al 2013)), then incubated for 2 to 3 h, shallowly submerged in buffered-HEPES, in a humidified chamber.

### Optophysiology, acquisition

GCaMP fluorescence fluctuations were imaged in an upright Thorlabs Bergamo II two-photon microscope (Thorlabs), with a 16x physiological objective (Nikon) and multiphoton excitation set at 920 nm. The light-path allowed a separation of red and green fluorescence at two concurrent photo-multiplier tubes (PMT), or green fluorescence with concurrent retinal photoreceptor stimulation with a lime LED (Thorlabs M565L3, 550-575, λmax 565 nm). During imaging, the retina was superfused with bicarbonate-buffered Ames solution that was constantly equilibrated with 95% O_2_ and 5% CO_2_ and temperature controlled at 32°C with an in-line heater (TC-334, Warner Instrument Corporation, USA) controlled by a Thermocontroller (TC-344B, Warner Instrument Corporation, USA). Each explant was imaged at the medio-lateral region, to promote consistency by avoiding the optic nerve head as well as the retinal periphery.

### Optophysiology, live z-Stack imaging

Image acquisitions were performed with ThorImage 3.2 (Thorlabs, USA) and saved as Bio-Format TIFF folders (Open Microscopy Environment). The piezo-driven microscope stage enabled acquisition at 1 µm spatial resolution in Z at a spatial resolution of 1024×1024 pixels, with unidirectional scanning and 5x frame averaging. Data were imported and processed in FIJI (NIH) before exporting as csv files for further processing using custom Matlab scripts.

### Optophysiology, dynamic signal acquisition

Image acquisitions were performed with ThorImage 3.2 (Thorlabs, USA) and saved as 16-bit unsigned TIFF-stacks. Single plane acquisition (spontaneous signals) was performed at a spatial resolution of 1024×1024, magnification of 16x, spatial resolution of 0.815 µm/pixel and a temporal resolution of 1.53 frames/s (7.65 fps with 5x frame averaging). Multi-plane acquisition (Fast-Z) was performed with a galvo-resonant scanner (8 kHz), with 5 to 7 planes spaced 20 µm from each other, at a spatial resolution of 512×512 pixels and magnifications of either 32x (1.63 pixels/µm) or 64x (3.26 pixels/µm) at a final temporal resolution of 1.975 frames per second.

### Optophysiology, light stimulation

Light-stimulation was controlled by ThorSync 3.2, which mediated illumination and extinction of the stimulation LED by means of TTL signals at a sampling frequency of 30 kHz. Three main light stimulation protocols were employed: five repeats of 1s ON/4 seconds OFF; 20 repeats of 500 ms ON/500 ms OFF; and five repeats of 10s ON/ 120s OFF. Light stimulation was initiated after a 30 second period of adaptation to the illumination provided by the multiphoton calcium imaging system. Irradiance at the retinal sample was measured with an ILT950 spectroradiometer (Information Light Technologies LLC), and derived using Govadorwskii nomograms (Allen et al., 2014; Govardovskii, Calvert, & Arshavsky, 2000). Isomerizations estimates for the stimulating LED were 6.16x 10^12^ (m-cone opsin), 7.27 10^10^ (s-cone opsin), 4.31 10^12^ (rod opsin), 1.7x 10^12^ (melanopsin) photons/cm^2^/s.

### Optophysiology, pharmacology

The pharmacological compounds (with final concentration and vendor) used in this study were AP4 (20 µM, Tocris Bioscience), AP5 (50 µM, Tocris Bioscience), CNQX (10 µM, Tocris Bioscience), Ouabain (10 µM, Sigma Aldrich) and TBOA (25 µM, Tocris Bioscience). These were diluted in bicarbonate-buffered Ames solution, constantly equilibrated with 95% O_2_ and 5% CO_2_. When studying glutamatergic blockade, each explant investigated on was subjected to all three blockers (CNQX, AP5 and AP4), but these were applied in a different order for each experiment. Pharmacological investigations consisted of the last step in optophysiological imaging for an explant, and the entire perfusion system was thoroughly cleaned with 70% ethanol and distilled water between explants.

### Exclusion criteria

For optophysiology, each retina yielded at least two explants, each of which underwent the entire acquisition protocol listed above. The data from some recordings were unusable due to recording artefacts, including too much mechanical drift, perfusion pump failure, microscope failure or light stimulation errors. The protocol was repeated if the failure was observed online. In the cases where it wasn’t, that particular recording was excluded, but other recordings from the same explant would be kept. As such, each experiment used a minimum N of 5 animals, although the number of explants is variable.

### Image processing for quantitative analyses of spontaneous calcium dynamics

Spontaneous calcium dynamics were studied on 11-minute duration recordings at a magnification of 16x, recorded in a single plane and sampled at 1.53 fps, with online averaging of 5 frames. Neural and vascular structures were readily identifiable from observations of every single frame, as in all ages and phenotypes they displayed strong resting fluorescence. Glial structures had to be searched for manually by observing every single frame in a recording, as they only appeared in bright, sporadic bursts. Five neural and vascular ROIs were chosen in random positions of the visual field and corresponded to 100×100 pixel squares or an ellipse with perimeter < 100 pixels, respectively. The raw fluorescence signal for each ROI was extracted in Fiji (NIH), then exported to Matlab as CSV files and processed with custom scripts. Fluorescent signals (F) were normalized (*Δ*F/F) by the following formula:

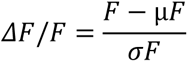

with µF and σF corresponding to the mean and standard deviation of the fluorescence over each pixel within an ROI and over the entire time series for each recording.

Waves and wavelets were thresholded with 4x standard deviations over a sliding average. Wavelet metrics (peak-to-trough amplitude, inter-wave-interval, wave duration) were exported and plotted as distributions or averaged by wave, ROI, explant, animal, for each condition using the formula below:

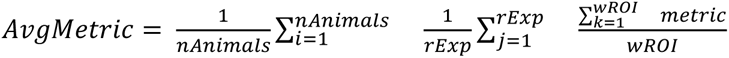

With nAnimals corresponding to the number of animals per condition, rExp corresponding to the number of ROIs per explant (5, for neuronal and vascular signals) and w corresponding to the number of waves detected in each ROI. Propagation distance and velocity were calculated manually in FiJi (NIH) for 5 events per explant per animal.

The correlation coefficient matrix for 5 neuronal and 5 vascular ROIs (**Figure S4, C**) was calculated in Matlab, before removing the diagonal auto-cross-correlation coefficients and averaging the neuro-neuro, neuro-vascular and vascular-vascular cross-correlation coefficients of those 5 ROIs for each explant (**Figure S4, D**).

### Image processing for pharmacological experiments

All pharmacological experiments were performed with 6-plane multilayered imaging, and two fly-back frames. For neuronal signals, an ROI of 100×100 was applied manually to an area within a neural field where wavelets were clearly observed. The raw fluorescence signal was extracted in Fiji (NIH), then exported to Matlab as CSV files and processed with custom scripts which detected wavelets using unbiased thresholding (sliding average + 4x sliding STD), then evaluated their frequency and amplitude in each retinal layer and under different pharmacological conditions. Glial events were evaluated in a similar manner, with the exception that a new ROI was generated manually for each glial event, and metrics included the surface of glial propagation.

### Image processing of multiphoton data for publication format

For videos, files were imported into Fiji (NIH), and then averaged in groups of 4 images along the temporal axis. Brightness and contrast were balanced manually to enhance calcium signals. Videos were saved as Jpeg-compressed FLV files, with a frame rate of 8 fps. This resulted in an effective frame rate of 24 fps, corresponding to an accelerated display speed of approximately 16x. For temporally projected still images, stacks were processed in Fiji (subtracted background, Gaussian blur) before manual thresholding. Further binary processing was necessary to highlight waves or wavelets before projection of the temporal hyperstack using Fiji’s “Fire” lookup table.

### Wholemount IHC

Labelling of glutamine synthetase (GS) in the retina highlights Muller glia, whilst glial fibrillary acidic protein (GFAP) highlights astrocytes and immune-responsive muller cells; a subset of RGCs express SMI-32. The primary antibodies used in this study were goat anti-mouse albumin (Bethyl, A90-234A, 1:800, RRID:AB_67122); mouse anti-GS (Millipore MAB302, 1:2000); chicken anti-GFAP (Chemicon, AB5541, 1:3000, RRID:AB_177521), mouse anti-SMI-32 (Covance, SMI-32R, 1:2000) and Hoechst 33342 (Sigma Aldrich, 1:300) for nuclear labeling. The secondary antibodies were conjugated to Alexa 488 (1:1000; green fluorescence, Molecular Probes), Cy3 (1:500 red fluorescence, Jackson ImmunoResearch) and Cy5 (1:500; far red fluorescence, Jackson ImmunoResearch). Following optophysiological experiments, explants were fixed in a solution of 4% Paraformaldehyde (PFA) in PBS for 15 minutes. After fixation, explants were washed three times for 15 minutes in PBS, then blocked for 10 h in CTA, a PBS solution containing 5% chemiblocker (membrane-blocking agent, Chemicon), 0.5% Triton X-100 and 0.05% sodium azide (Sigma).

Primary antibodies were diluted in CTA and incubated for 72 h, followed by incubation for 48 h in the appropriate secondary antibody and nuclear stain. After staining, explants were flat-mounted on a slide, GCL up, and coverslipped using Vectashield mounting medium (H-1000, Vector Laboratories). The coverslip was sealed in place with transparent nail varnish. To avoid compression of retinal explants, small pieces of broken glass coverslips (1mm thick) were placed in the space between the slide and the coverslip, on either side of each explant. All steps were completed at room temperature, except for incubation of the GS primary antibody, which required a 37° C incubation while being gently shaken. Slides were imaged in an upright Leica SP8 or inverted Nikon A1R HD25 confocal microscope. Data were imported and processed in FIJI (NIH).

### Statistics

Statistical analyses and graph generation were performed in Prism (GraphPad) version 8.2 and version 9.1.0 for RS1 expression analyses. Mann-Whitney test was used for comparing non-parametric datasets in groups of 2. The Kruskall-Wallis test with Dunn’s multiple comparisons was used to compare groups of 3 or more non-parametric datasets. Quantifications of neuro-vascular correlations analyses were analyzed using two-way ANOVA with Šidák’s multiple comparisons test. *Rs1* expression analyses were analyzed using one-way ANOVA for each retinal layer with Dunnett’s multiple comparisons test.

## Author Contributions

BS, NSP and CE designed the study. CE and CC performed optophysiology experiments. CE, SK, YL and DS performed histology experiments. RS and JB performed OCT experiments. CE and SK analyzed the data. CE wrote the manuscript with extensive contributions from the other authors.

## Conflict of Interest Statement

Shireen Khattak, Elena Ivanova, Yang Liu, Duo Sun, Carmelo Romano and Botir Sagdullaev are employees of Regeneron Pharmaceuticals.

## Acknowledgements

We thank Susan Croll for helpful discussions on biostatistics and Lampros Panagis for help with setting up imaging of retinoschisin protein and RNA.

## Funding

This work was supported by grants from the National Institutes of Health [R01 EY029796 to N.S.P. J.M. & B.T.S.; R01 EY026576 to B.T.S.] and by core grants to the Departments of Ophthalmology of the Cleveland Clinic Lerner College of Medicine of CWRU (CCLCM of CWRU; P30 EY025585), and the University of Illinois at Chicago College of Medicine (UIC COM; P30 EY001792); by unrestricted grants from Research to Prevent Blindness to the Departments of Ophthalmology of CCLCM of CWRU and UIC COM.; and by Regeneron Pharmaceuticals. N.S.P. is a VA Research Career Scientist.

## Supplementary Figures

**Figure S1:**
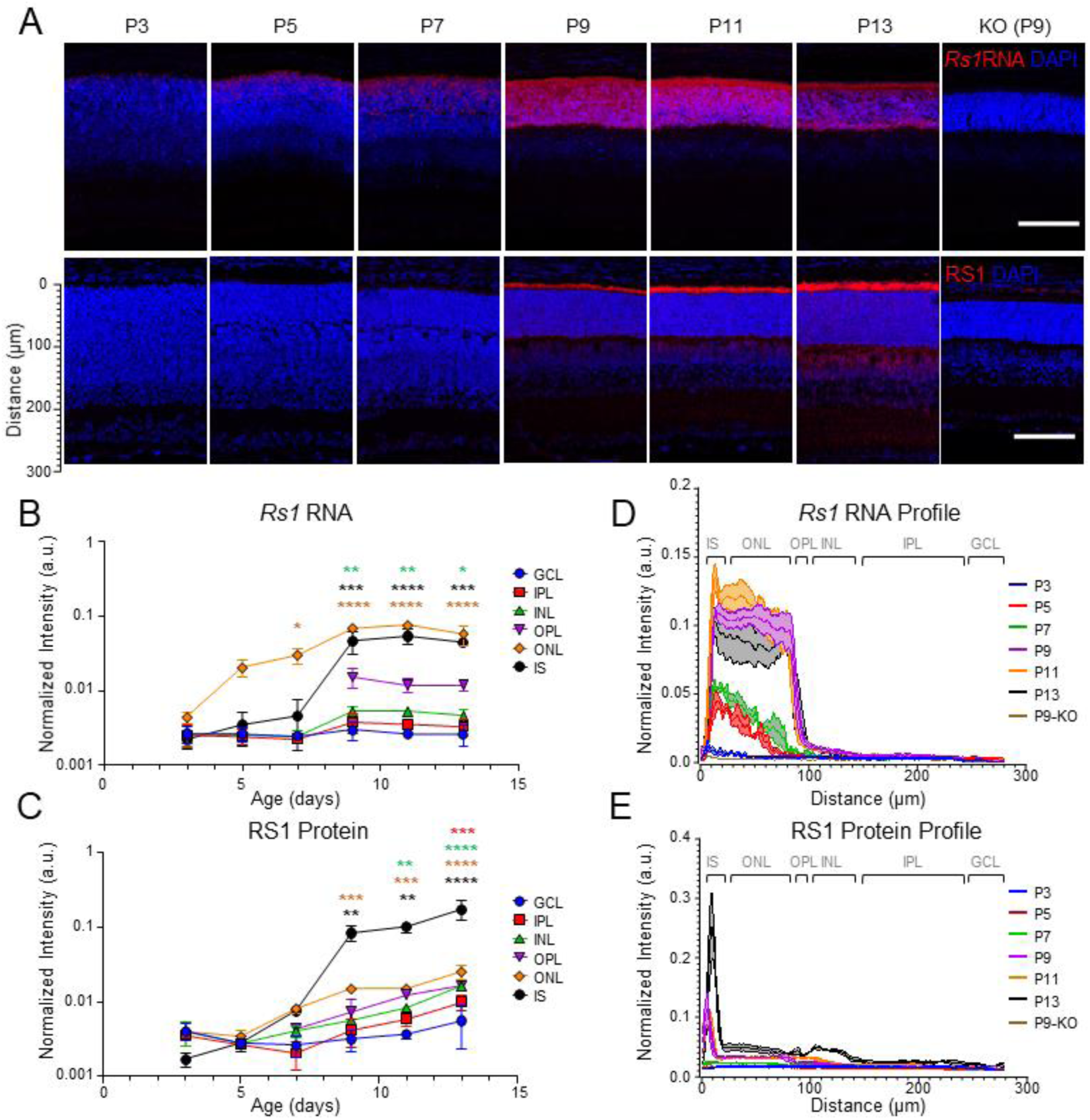
RS1 is expressed throughout the retina during development. **A.** Retinal sections stained for nuclei (DAPI, blue), *Rs1* RNA (red, top) and RS1 protein (red, bottom). Scale bar is 100 µm. **B-C.** Fluorescence intensity in the red channel for *Rs1* RNA **(B)** and RS1 protein **(C)** in different layers of the retina. One-way ANOVA with multiple comparisons for each retinal layer compared to P3, N = 3 mice per condition. For **(B)** p = 0.0023, 0.0024 & <0.0001 (INL at P9, P11 and P13, respectively); p = 0.0195 at P7, & < 0.0001 at P9, P11 and P13 (ONL); p = 0.0002, < 0.0001 & =0.0003 (IS at P9, P11 and P13, respectively). For **(C)** p = 0.003 (IPL); 0.0011 & <0.0001 (INL at P11 and P13, respectively); p = 0.0008, 0.0008 & <0.0001 (ONL at P9, P11 and P13, respectively); p = 0.0052, 0.0011 & <0.0001 (IS at P9, P11 and P13, respectively). **D-E** Normalized mean fluorescence (±SEM) intensity profile through retinal cross sections from the edge of photoreceptor inner segments in the red channel for *Rs1* RNA **(D)** and RS1 protein **(E)**. Distance scale diagrammed in **(A)**. 16 bit images with max grey value of 65,535. GCL: Ganglion Cell Layer; IPL: Inner Plexiform Layer; INL: Inner Nuclear Layer; OPL: Outer Plexiform Layer; ONL: Outer Nuclear Layer; IS: Inner Segments.

**Figure S2:**
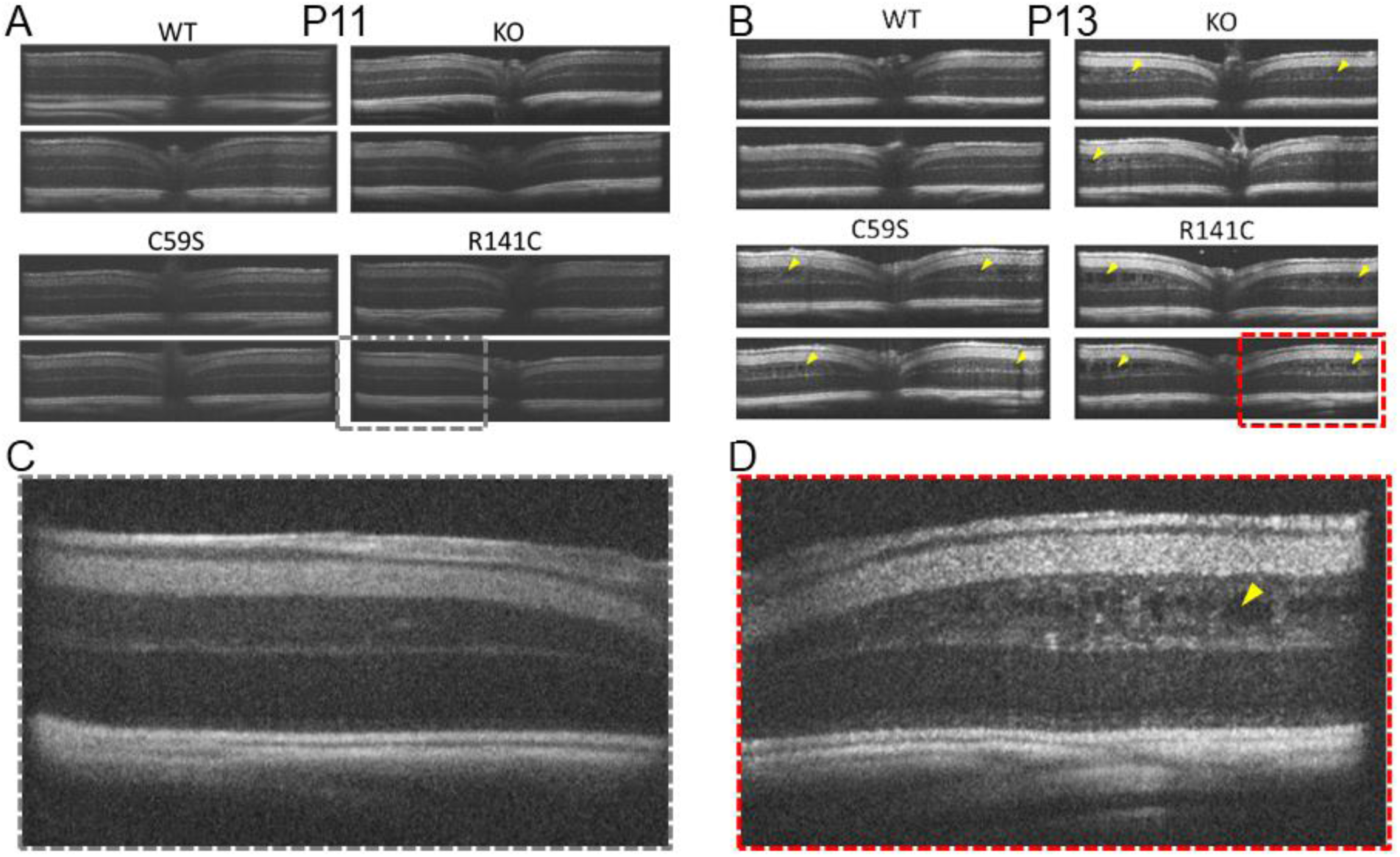
Schisis appears at P13 in 3 mouse models of XLRS. **A-B.** OCT imaging of mouse models at P 11 **(A)** and 13 **(B)**. **A.** At P11, the XLRS models are indistinguishable from the control retina. **B.** At P13, all three XLRS models display schisis (yellow arrowheads). **C-D.** Zoom of half portion of R141C retina at P11 **(C)** and P13 **(D)**. Images are representative of 5-6 mice for each genotype and age.

**Figure S3:**
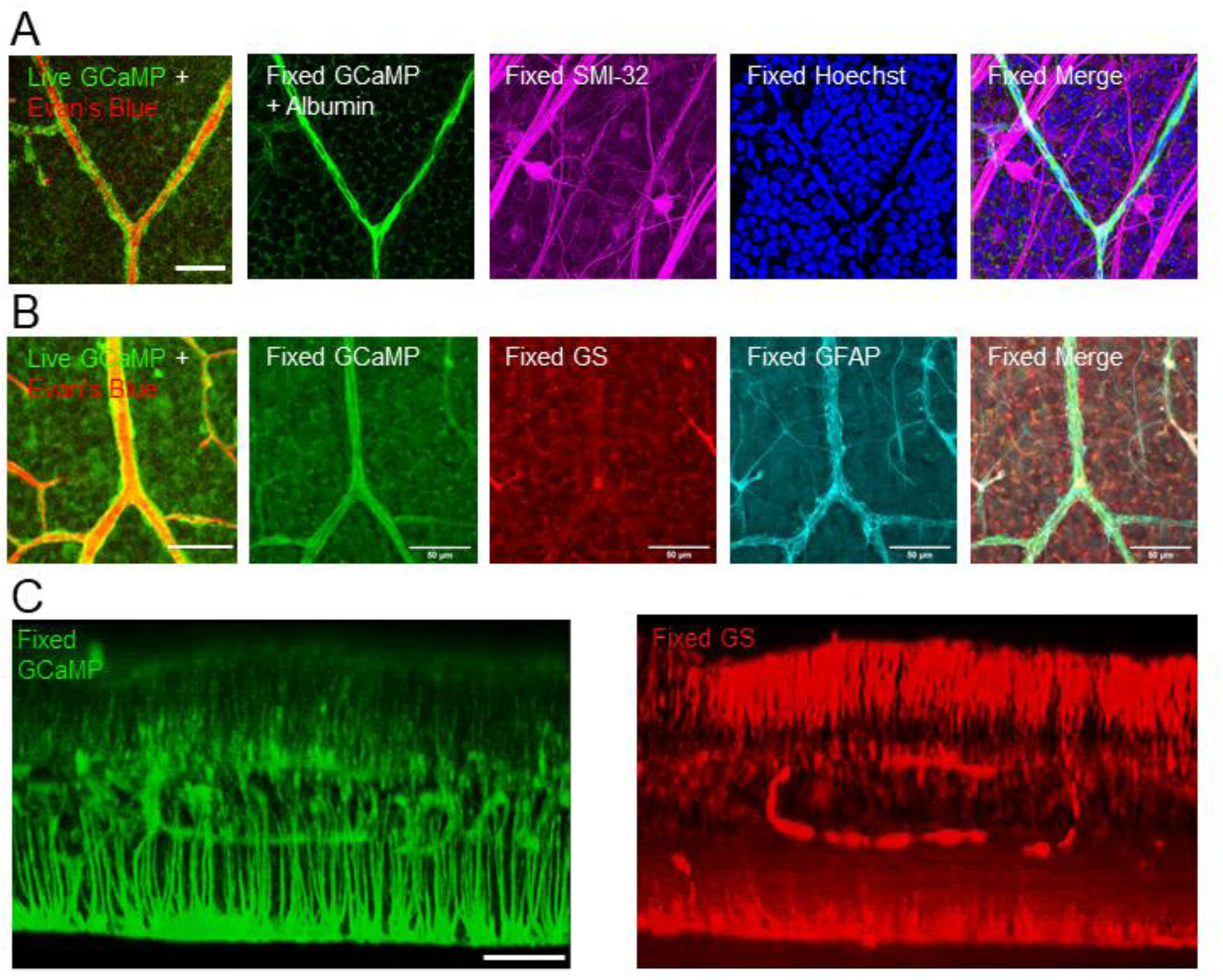
GCaMP6f is expressed in neurons, glia and vascular cells in the NG2 GCAMP mouse line. **A.** Live mature retinal tissue compared to stain for neurons (SMI-32) vascular cells (Albumin) and nuclear DNA (Hoechst). **B.** Live retinal tissue compared to stain for glia. Glutamine synthetase (GS) is specific to Müller glia, but glial fibrillary acidic protein (GFAP) is expressed in both Müller glia and astrocytes. **C.** 3D projected retinal section, with interpolation, displaying how GCaMP is highly expressed in Muller cells.

**Figure S4:**
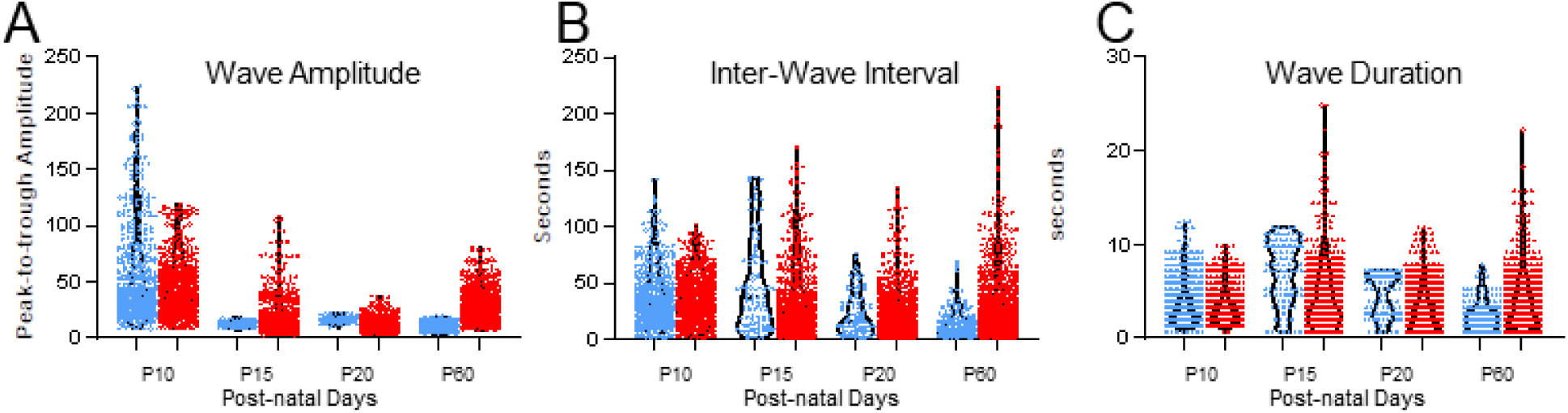
Metrics for each individually thresholded neuronal events. Violin-plot distributions of neural wavelet Amplitude (**A**), Inter-wave-interval (**B**), Duration (**C**), displaying individual values for each wavelet detected.

**Figure S5:**
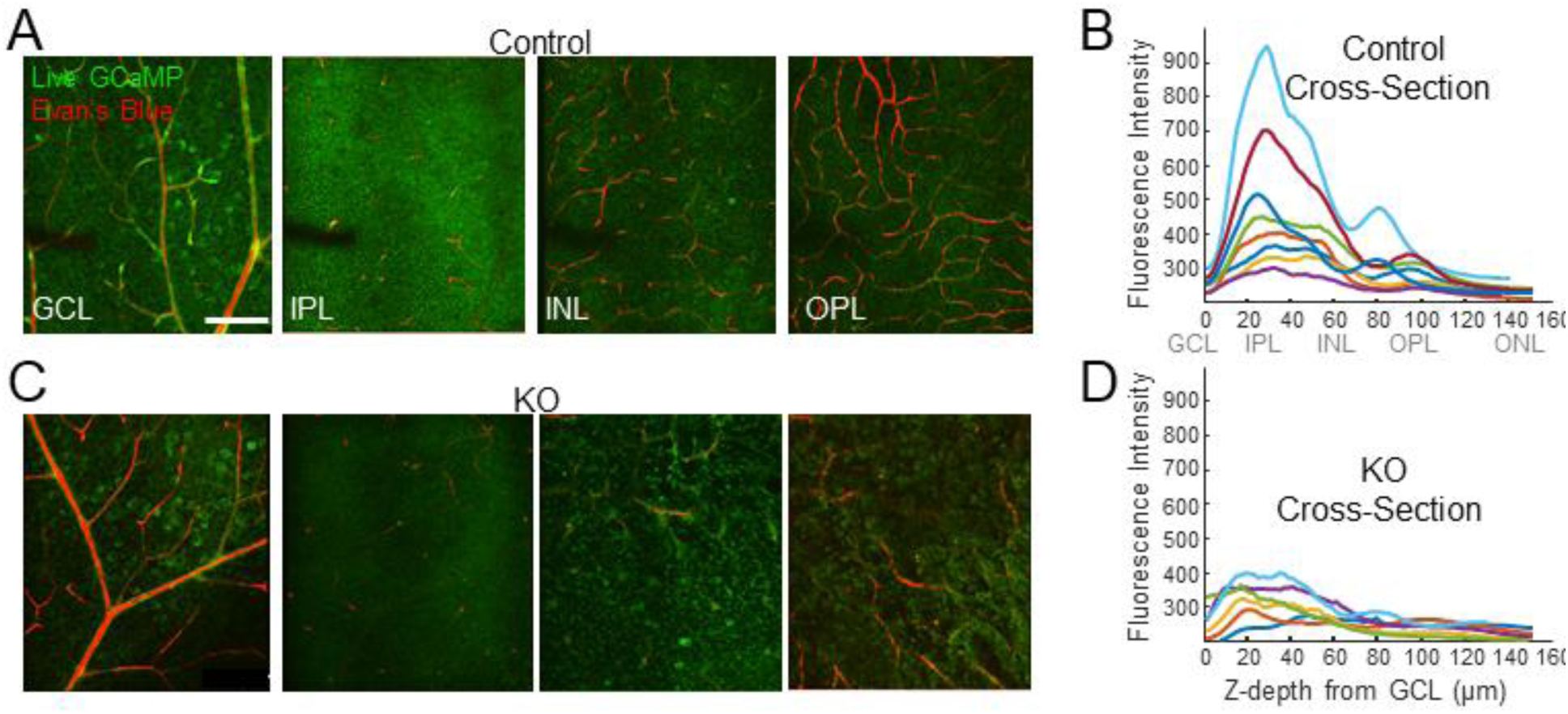
Disrupted landscape of RS1-deficient retina. **A.** Live control retina displays typical lamination profile, with blood vessels marking GCL, INL and OPL. **B.** The Z-profile of GCaMP6f fluorescence from 6 control retinas displays a bimodal distribution, with peaks in the synaptic layers. **C.** KO retinas display fibrous appearance in non-neuronal layers. **(D)** The Z-profile of GCaMP6f fluorescence from 6 KO retinas displays a stunted distribution, with no clear peaks. The grey legend between B and D provides a rough guide to retinal layers according to depth from GCL surface.

## References

Alarcon-Martinez, L., Villafranca-Baughman, D., Quintero, H., Kacerovsky, J. B., Dotigny, F., Murai, K. K., … Di Polo, A. (2020). Interpericyte tunnelling nanotubes regulate neurovascular coupling. Nature, 585(7823), 91–95. doi:10.1038/s41586-020-2589-x

Alexander, K. R., Barnes, C. S., & Fishman, G. A. (2005). Characteristics of contrast processing deficits in X-linked retinoschisis. Vision Res, 45(16), 2095–2107. doi:10.1016/j.visres.2005.01.037

Allen, A. E., Storchi, R., Martial, F. P., Petersen, R. S., Montemurro, M. A., Brown, T. M., & Lucas, R. J. (2014). Melanopsin-driven light adaptation in mouse vision. Curr Biol, 24(21), 2481–2490. doi:10.1016/j.cub.2014.09.015

Ambrosio, L., Williams, J. S., Gutierrez, A., Swanson, E. A., Munro, R. J., Ferguson, R. D., … Akula, J. D. (2021). Carbonic anhydrase inhibition in X-linked retinoschisis: An eye on the photoreceptors. Exp Eye Res, 202, 108344. doi:10.1016/j.exer.2020.108344

Baden, T., Berens, P., Franke, K., Roman Roson, M., Bethge, M., & Euler, T. (2016). The functional diversity of retinal ganglion cells in the mouse. Nature, 529(7586), 345–350. doi:10.1038/nature16468

Baden, T., Euler, T., & Berens, P. (2020). Understanding the retinal basis of vision across species. Nat Rev Neurosci, 21(1), 5–20. doi:10.1038/s41583-019-0242-1

Bansal, A., Singer, J. H., Hwang, B. J., Xu, W., Beaudet, A., & Feller, M. B. (2000). Mice lacking specific nicotinic acetylcholine receptor subunits exhibit dramatically altered spontaneous activity patterns and reveal a limited role for retinal waves in forming ON and OFF circuits in the inner retina. J Neurosci, 20(20), 7672–7681.

Bell, B. A., Kaul, C., Bonilha, V. L., Rayborn, M. E., Shadrach, K., & Hollyfield, J. G. (2015). The BALB/c mouse: Effect of standard vivarium lighting on retinal pathology during aging. Exp Eye Res, 135, 192–205. doi:10.1016/j.exer.2015.04.009

Bell, B. A., Kaul, C., & Hollyfield, J. G. (2014). A protective eye shield for prevention of media opacities during small animal ocular imaging. Exp Eye Res, 127, 280–287. doi:10.1016/j.exer.2014.01.001

Bringmann, A., Pannicke, T., Biedermann, B., Francke, M., Iandiev, I., Grosche, J., … Reichenbach, A. (2009). Role of retinal glial cells in neurotransmitter uptake and metabolism. Neurochem Int, 54(3-4), 143–160. doi:10.1016/j.neuint.2008.10.014

Bush, M., Setiaputra, D., Yip, C. K., & Molday, R. S. (2016). Cog-Wheel Octameric Structure of RS1, the Discoidin Domain Containing Retinal Protein Associated with X-Linked Retinoschisis. PLoS One, 11(1), e0147653. doi:10.1371/journal.pone.0147653

Chen, D., Xu, T., Tu, M., Xu, J., Zhou, C., Cheng, L., … Gu, F. (2017). Recapitulating X-Linked Juvenile Retinoschisis in Mouse Model by Knock-In Patient-Specific Novel Mutation. Front Mol Neurosci, 10, 453. doi:10.3389/fnmol.2017.00453

Chen, T. W., Wardill, T. J., Sun, Y., Pulver, S. R., Renninger, S. L., Baohan, A., … Kim, D. S. (2013). Ultrasensitive fluorescent proteins for imaging neuronal activity. Nature, 499(7458), 295–300. doi:10.1038/nature12354

Cukras, C., Wiley, H. E., Jeffrey, B. G., Sen, H. N., Turriff, A., Zeng, Y., … Sieving, P. A. (2018). Retinal AAV8-RS1 Gene Therapy for X-Linked Retinoschisis: Initial Findings from a Phase I/IIa Trial by Intravitreal Delivery. Mol Ther, 26(9), 2282–2294. doi:10.1016/j.ymthe.2018.05.025

Dalkara, D., Byrne, L. C., Klimczak, R. R., Visel, M., Yin, L., Merigan, W. H., … Schaffer, D. V. (2013). In vivo-directed evolution of a new adeno-associated virus for therapeutic outer retinal gene delivery from the vitreous. Sci Transl Med, 5(189), 189ra176. doi:10.1126/scitranslmed.3005708

Demas, J., Sagdullaev, B. T., Green, E., Jaubert-Miazza, L., McCall, M. A., Gregg, R. G., … Guido, W. (2006). Failure to maintain eye-specific segregation in nob, a mutant with abnormally patterned retinal activity. Neuron, 50(2), 247–259. doi:10.1016/j.neuron.2006.03.033

Derouiche, A., & Rauen, T. (1995). Coincidence of L-glutamate/L-aspartate transporter (GLAST) and glutamine synthetase (GS) immunoreactions in retinal glia: evidence for coupling of GLAST and GS in transmitter clearance. J Neurosci Res, 42(1), 131–143. doi:10.1002/jnr.490420115

Diamond, J. S., & Copenhagen, D. R. (1993). The contribution of NMDA and non-NMDA receptors to the light-evoked input-output characteristics of retinal ganglion cells. Neuron, 11(4), 725–738. doi:10.1016/0896-6273(93)90082-3

Feller, M. B. (2009). Retinal waves are likely to instruct the formation of eye-specific retinogeniculate projections. Neural Dev, 4, 24. doi:10.1186/1749-8104-4-24

Feller, M. B., Wellis, D. P., Stellwagen, D., Werblin, F. S., & Shatz, C. J. (1996). Requirement for cholinergic synaptic transmission in the propagation of spontaneous retinal waves. Science, 272(5265), 1182–1187. doi:10.1126/science.272.5265.1182

Forsius, H., Krause, U., Helve, J., Vuopala, V., Mustonen, E., Vainio-Mattila, B., … Eriksson, A. W. (1973). Visual acuity in 183 cases of X-chromosomal retinoschisis. Can J Ophthalmol, 8(3), 385–393.

Franke, K., Berens, P., Schubert, T., Bethge, M., Euler, T., & Baden, T. (2017). Inhibition decorrelates visual feature representations in the inner retina. Nature, 542(7642), 439–444. doi:10.1038/nature21394

Friedrich, U., Stohr, H., Hilfinger, D., Loenhardt, T., Schachner, M., Langmann, T., & Weber, B. H. (2011). The Na/K-ATPase is obligatory for membrane anchorage of retinoschisin, the protein involved in the pathogenesis of X-linked juvenile retinoschisis. Hum Mol Genet, 20(6), 1132–1142. doi:10.1093/hmg/ddq557

Gargini, C., Terzibasi, E., Mazzoni, F., & Strettoi, E. (2007). Retinal organization in the retinal degeneration 10 (rd10) mutant mouse: a morphological and ERG study. J Comp Neurol, 500(2), 222–238. doi:10.1002/cne.21144

George, N. D., Yates, J. R., & Moore, A. T. (1995). X linked retinoschisis. Br J Ophthalmol, 79(7), 697–702. doi:10.1136/bjo.79.7.697

Govardovskii, V. I., Calvert, P. D., & Arshavsky, V. Y. (2000). Photoreceptor light adaptation. Untangling desensitization and sensitization. J Gen Physiol, 116(6), 791–794. doi:10.1085/jgp.116.6.791

Hippert, C., Graca, A. B., Barber, A. C., West, E. L., Smith, A. J., Ali, R. R., & Pearson, R. A. (2015). Muller glia activation in response to inherited retinal degeneration is highly varied and disease-specific. PLoS One, 10(3), e0120415. doi:10.1371/journal.pone.0120415

Jablonski, M. M., Dalke, C., Wang, X., Lu, L., Manly, K. F., Pretsch, W., … Graw, J. (2005). An ENU-induced mutation in Rs1h causes disruption of retinal structure and function. Mol Vis, 11, 569–581.

Jones, B. W., Kondo, M., Terasaki, H., Lin, Y., McCall, M., & Marc, R. E. (2012). Retinal remodeling. Jpn J Ophthalmol, 56(4), 289–306. doi:10.1007/s10384-012-0147-2

Jones, B. W., Watt, C. B., & Marc, R. E. (2005). Retinal remodelling. Clin Exp Optom, 88(5), 282–291. doi:10.1111/j.1444-0938.2005.tb06712.x

Kalloniatis, M., Sun, D., Foster, L., Haverkamp, S., & Wassle, H. (2004). Localization of NMDA receptor subunits and mapping NMDA drive within the mammalian retina. Vis Neurosci, 21(4), 587–597. doi:10.1017/s0952523804214080

Kovacs-Oller, T., Ivanova, E., Bianchimano, P., & Sagdullaev, B. T. (2020). The pericyte connectome: spatial precision of neurovascular coupling is driven by selective connectivity maps of pericytes and endothelial cells and is disrupted in diabetes. Cell Discov, 6, 39. doi:10.1038/s41421-020-0180-0

Liu, Y., Kinoshita, J., Ivanova, E., Sun, D., Li, H., Liao, T., … Romano, C. (2019). Mouse models of X-linked juvenile retinoschisis have an early onset phenotype, the severity of which varies with genotype. Hum Mol Genet, 28(18), 3072–3090. doi:10.1093/hmg/ddz122

Maccione, A., Hennig, M. H., Gandolfo, M., Muthmann, O., van Coppenhagen, J., Eglen, S. J., … Sernagor, E. (2014). Following the ontogeny of retinal waves: pan-retinal recordings of population dynamics in the neonatal mouse. J Physiol, 592(7), 1545–1563. doi:10.1113/jphysiol.2013.262840

Meister, M., Wong, R. O., Baylor, D. A., & Shatz, C. J. (1991). Synchronous bursts of action potentials in ganglion cells of the developing mammalian retina. Science, 252(5008), 939–943. doi:10.1126/science.2035024

Mishra, A., Vijayasarathy, C., Cukras, C. A., Wiley, H. E., Sen, H. N., Zeng, Y., … Sieving, P. A. (2021). Immune function in X-linked retinoschisis subjects in an AAV8-RS1 phase I/IIa gene therapy trial. Mol Ther, 29(6), 2030–2040. doi:10.1016/j.ymthe.2021.02.013

Molday, L. L., Min, S. H., Seeliger, M. W., Wu, W. W., Dinculescu, A., Timmers, A. M., … Molday, R. S. (2006). Disease mechanisms and gene therapy in a mouse model for X-linked retinoschisis. Adv Exp Med Biol, 572, 283–289. doi:10.1007/0-387-32442-9_39

Molday, L. L., Wu, W. W., & Molday, R. S. (2007). Retinoschisin (RS1), the protein encoded by the X-linked retinoschisis gene, is anchored to the surface of retinal photoreceptor and bipolar cells through its interactions with a Na/K ATPase-SARM1 complex. J Biol Chem, 282(45), 32792–32801. doi:10.1074/jbc.M706321200

Molday, R. S., Kellner, U., & Weber, B. H. (2012). X-linked juvenile retinoschisis: clinical diagnosis, genetic analysis, and molecular mechanisms. Prog Retin Eye Res, 31(3), 195–212. doi:10.1016/j.preteyeres.2011.12.002

Nakajima, Y., Iwakabe, H., Akazawa, C., Nawa, H., Shigemoto, R., Mizuno, N., & Nakanishi, S. (1993). Molecular characterization of a novel retinal metabotropic glutamate receptor mGluR6 with a high agonist selectivity for L-2-amino-4-phosphonobutyrate. J Biol Chem, 268(16), 11868–11873.

Ou, J., Vijayasarathy, C., Ziccardi, L., Chen, S., Zeng, Y., Marangoni, D., … Sieving, P. A. (2015). Synaptic pathology and therapeutic repair in adult retinoschisis mouse by AAV-RS1 transfer. J Clin Invest, 125(7), 2891–2903. doi:10.1172/JCI81380

Peachey, N. S., Fishman, G. A., Derlacki, D. J., & Brigell, M. G. (1987). Psychophysical and electroretinographic findings in X-linked juvenile retinoschisis. Arch Ophthalmol, 105(4), 513–516. doi:10.1001/archopht.1987.01060040083038

Pennesi, M. E., Birch, D. G., Jayasundera, K. T., Parker, M., Tan, O., Gurses-Ozden, R., … For The Xlrs-Study, G. (2018). Prospective Evaluation of Patients With X-Linked Retinoschisis During 18 Months. Invest Ophthalmol Vis Sci, 59(15), 5941–5956. doi:10.1167/iovs.18-24565

Plossl, K., Royer, M., Bernklau, S., Tavraz, N. N., Friedrich, T., Wild, J., … Friedrich, U. (2017). Retinoschisin is linked to retinal Na/K-ATPase signaling and localization. Mol Biol Cell, 28(16), 2178–2189. doi:10.1091/mbc.E17-01-0064

Plossl, K., Weber, B. H., & Friedrich, U. (2017). The X-linked juvenile retinoschisis protein retinoschisin is a novel regulator of mitogen-activated protein kinase signalling and apoptosis in the retina. J Cell Mol Med, 21(4), 768–780. doi:10.1111/jcmm.13019

Rosa, J. M., Bos, R., Sack, G. S., Fortuny, C., Agarwal, A., Bergles, D. E., … Feller, M. B. (2015). Neuron-glia signaling in developing retina mediated by neurotransmitter spillover. Elife, 4. doi:10.7554/eLife.09590

Sauer, C. G., Gehrig, A., Warneke-Wittstock, R., Marquardt, A., Ewing, C. C., Gibson, A., … Weber, B. H. (1997). Positional cloning of the gene associated with X-linked juvenile retinoschisis. Nat Genet, 17(2), 164–170. doi:10.1038/ng1097-164

Takada, Y., Fariss, R. N., Tanikawa, A., Zeng, Y., Carper, D., Bush, R., & Sieving, P. A. (2004). A retinal neuronal developmental wave of retinoschisin expression begins in ganglion cells during layer formation. Invest Ophthalmol Vis Sci, 45(9), 3302–3312. doi:10.1167/iovs.04-0156

Takada, Y., Vijayasarathy, C., Zeng, Y., Kjellstrom, S., Bush, R. A., & Sieving, P. A. (2008). Synaptic pathology in retinoschisis knockout (Rs1-/y) mouse retina and modification by rAAV-Rs1 gene delivery. Invest Ophthalmol Vis Sci, 49(8), 3677–3686. doi:10.1167/iovs.07-1071

Tanino, T., Katsumi, O., & Hirose, T. (1985). Electrophysiological similarities between two eyes with X-linked recessive retinoschisis. Doc Ophthalmol, 60(2), 149–161. doi:10.1007/BF00158030

Testa, F., Di Iorio, V., Gallo, B., Marchese, M., Nesti, A., De Rosa, G., … Simonelli, F. (2019). Carbonic anhydrase inhibitors in patients with X-linked retinoschisis: effects on macular morphology and function. Ophthalmic Genet, 40(3), 207–212. doi:10.1080/13816810.2019.1616303

Thobani, A., & Fishman, G. A. (2011). The use of carbonic anhydrase inhibitors in the retreatment of cystic macular lesions in retinitis pigmentosa and X-linked retinoschisis. Retina, 31(2), 312–315. doi:10.1097/IAE.0b013e3181e587f9

Tolun, G., Vijayasarathy, C., Huang, R., Zeng, Y., Li, Y., Steven, A. C., … Heymann, J. B. (2016). Paired octamer rings of retinoschisin suggest a junctional model for cell-cell adhesion in the retina. Proc Natl Acad Sci U S A, 113(19), 5287–5292. doi:10.1073/pnas.1519048113

Weber, B. H., Schrewe, H., Molday, L. L., Gehrig, A., White, K. L., Seeliger, M. W., … Molday, R. S. (2002). Inactivation of the murine X-linked juvenile retinoschisis gene, Rs1h, suggests a role of retinoschisin in retinal cell layer organization and synaptic structure. Proc Natl Acad Sci U S A, 99(9), 6222–6227. doi:10.1073/pnas.092528599

Wu, W. W., Wong, J. P., Kast, J., & Molday, R. S. (2005). RS1, a discoidin domain-containing retinal cell adhesion protein associated with X-linked retinoschisis, exists as a novel disulfide-linked octamer. J Biol Chem, 280(11), 10721–10730. doi:10.1074/jbc.M413117200

Young, R. W. (1985). Cell differentiation in the retina of the mouse. Anat Rec, 212(2), 199–205. doi:10.1002/ar.1092120215

Yu, D. Y., Cringle, S., Valter, K., Walsh, N., Lee, D., & Stone, J. (2004). Photoreceptor death, trophic factor expression, retinal oxygen status, and photoreceptor function in the P23H rat. Invest Ophthalmol Vis Sci, 45(6), 2013–2019. doi:10.1167/iovs.03-0845

Zeng, Y., Takada, Y., Kjellstrom, S., Hiriyanna, K., Tanikawa, A., Wawrousek, E., … Sieving, P. A. (2004). RS-1 Gene Delivery to an Adult Rs1h Knockout Mouse Model Restores ERG b-Wave with Reversal of the Electronegative Waveform of X-Linked Retinoschisis. Invest Ophthalmol Vis Sci, 45(9), 3279–3285. doi:10.1167/iovs.04-0576

Zhu, X., Bergles, D. E., & Nishiyama, A. (2008). NG2 cells generate both oligodendrocytes and gray matter astrocytes. Development, 135(1), 145–157. doi:10.1242/dev.004895

